# Integrated Genomic Analysis Reveals Key Features of Long Undecoded Transcript Isoform (LUTI)-based Gene Repression

**DOI:** 10.1101/843458

**Authors:** Amy Tresenrider, Victoria Jorgensen, Minghao Chia, Hanna Liao, Folkert J. van Werven, Elçin Ünal

## Abstract

<u>L</u>ong <u>U</u>ndecoded <u>T</u>ranscript Isoforms (LUTIs) represent a class of non-canonical mRNAs that downregulate gene expression through the combined act of transcriptional and translational repression. While single gene studies revealed some important aspects of LUTI-based repression, how these features impact gene regulation at a global scale is unknown. By using transcript leader and direct RNA sequencing, here we identify 74 LUTI candidates that are expressed specifically during meiotic prophase. Translational repression of these candidates is ubiquitous and dependent on upstream open reading frames. However, LUTI-based transcriptional repression is highly variable. In only 50% of the cases, LUTI transcription causes downregulation of the protein-coding transcript isoform. Higher LUTI expression, enrichment of histone 3 lysine 36 trimethylation, and changes in nucleosome position are the strongest predictors of LUTI-based transcriptional repression. We conclude that LUTIs downregulate gene expression in a manner that integrates translational repression, chromatin state changes, and the magnitude of LUTI expression.

## INTRODUCTION

Gene expression programs determine cellular identity, form and function through extensive regulation at multiple levels. Although cells are equipped with a variety of machineries to control gene expression, transcription factors are thought to be the most critical regulators of cellular state change. Transcription factors are among the first genes to be expressed at the onset of a developmental program, where they initiate cascades of gene expression that ultimately drive cellular differentiation. While transcription factors are often implicated in gene activation, less focus is placed on how gene repression events can be coordinated with the transcription factor-dependent waves of gene activation.

New insights into a mechanism of transcription factor-mediated gene repression came from studying transcription and translation together, with the goal of elucidating the regulation of an essential gene called *NDC80*. Ndc80 is a conserved subunit of the kinetochore, the machinery that connects chromosomes to microtubules (reviewed in Ciferri et al., 2007; Tooley and Stukenberg, 2011). *NDC80* expression is tightly regulated during meiotic differentiation, the developmental pathway that generates reproductive cells (Asakawa et al., 2005; Chen et al., 2017; Miller et al., 2012; Sun et al., 2011). In budding yeast, regulation of *NDC80* is wired into the meiotic transcriptional program through the timely expression of two distinct mRNA isoforms. The promoters that drive the expression of these two mRNAs are located in tandem (Figure 1A). As cells enter meiosis, a key transcription factor complex composed of Ime1 and Ume6 activates the distal *NDC80* promoter (P2) to induce the expression of a 5’-extended mRNA isoform (Figure 1A, right panel). This transcript cannot be translated into a functional protein because the upstream open reading frames (uORFS) in its 5’-leader inhibit ORF translation by competitively engaging with the ribosomes. Instead of producing Ndc80 protein, the <u>L</u>ong <u>U</u>ndecoded <u>T</u>ranscript Isoform (LUTI) serves to inactivate the ORF-proximal *NDC80* promoter (P1) through co-transcriptional histone modifications. As a result, the level of the protein-coding *NDC80* transcript is dramatically reduced in meiotic prophase, leading to kinetochore inactivation (Chen et al., 2017; Chia et al., 2017). Prior to the meiotic divisions, the kinetochore complex is re-activated as a second meiotic transcription factor binds to the proximal *NDC80* promoter to produce a protein-coding transcript isoform. Accordingly, the developmental toggling between two functionally distinct *NDC80* mRNAs modulates the level of this essential kinetochore subunit during meiotic differentiation. Furthermore, since the LUTI-based repression of *NDC80* is intimately dependent on Ime1-Ume6, this mechanism provides an efficient strategy to coordinate gene activation and repression events through a common transcription factor.

**Figure 1.**
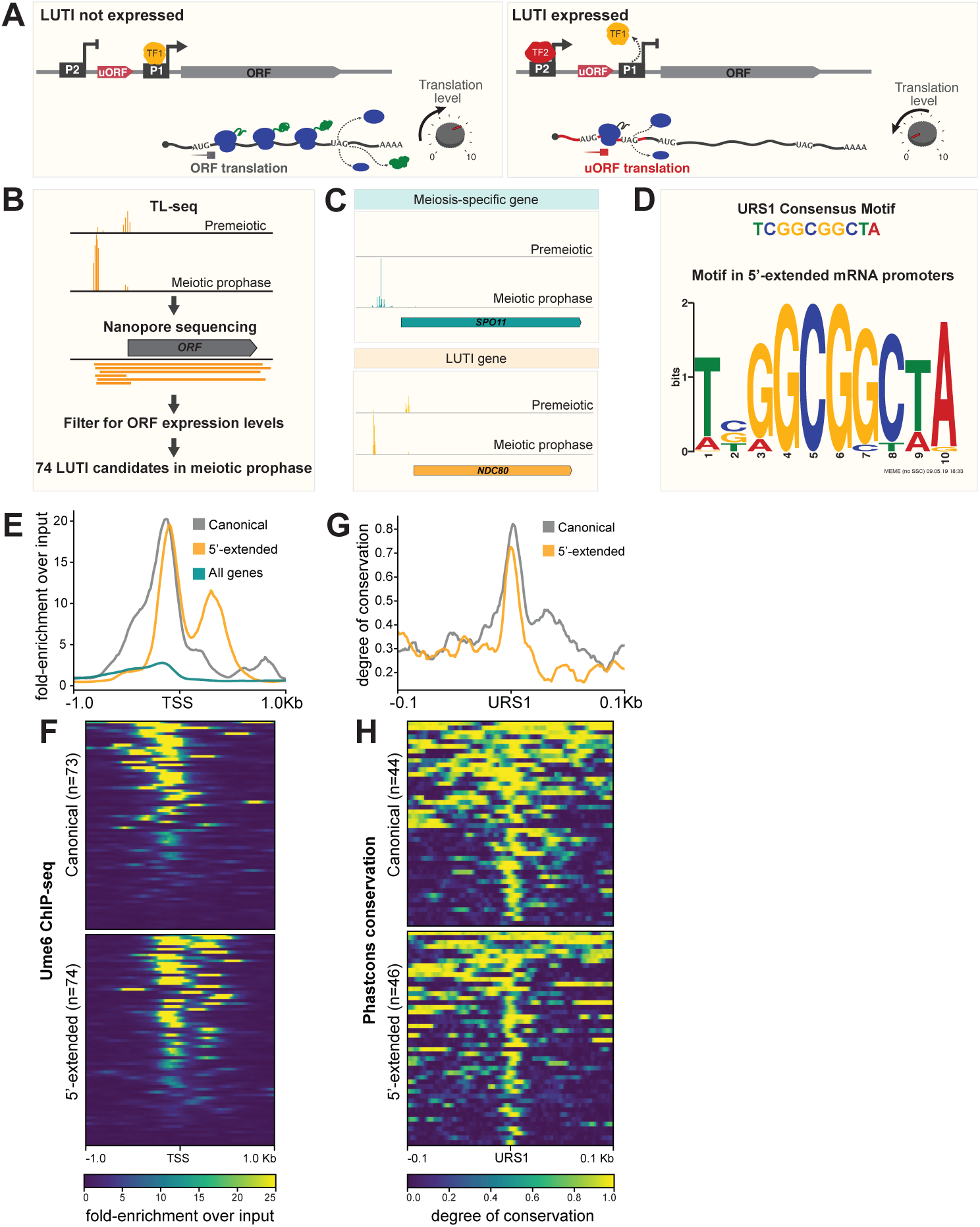
The pipeline for LUTI discovery and analysis of transcription-factor based regulation of LUTIs. **A.** A schematic of LUTI-based gene regulation. On the left is a depiction of a gene when the LUTI is not expressed and protein production is high. The right depicts features of LUTIs that lead to repression of protein production. **B.** An overview of the pipeline used to discover 5’-extended transcripts. Transcription starts sites (TSSs) that increase by at least a fold change of > 4 were identified by transcript leader sequencing (TL-seq). Long read nanopore sequencing was used to confirm that transcripts produced from these loci spanned the entire open reading frame of the downstream gene. Instances in which no ORF-proximal TSS was identified were not included in the analysis. **C.** Genome browser views of TL-seq for the meiosis-specific gene *SPO11* and the *NDC80* locus, which contains a LUTI. Note that *SPO11* is on the reverse strand, but the image has been flipped for clarity. **D.** The URS1 motif found in LUTI promoters. The consensus-binding motif observed in the 300 bp +/- of the distal TSSs from (B) as identified by MEME. In 50/74 instances, a significant URS1 binding motif match (combined match p-value < 0.05) was found. **E-F.** Ume6 ChIP-seq was performed in a strain with 3V5-tagged Ume6 (*UB3301*) grown in BYTA to saturation. **E.** Metagene analysis of Ume6 fold enrichment over input in the promoters of all genes compared to the promoters of previously identified Ume6 targets (Williams et al., 2002) and to the promoters of the 5’-extended transcripts identified in this study. A representative image from one of three replicates is shown. **F.** Heatmap of Ume6 fold enrichment over input in the promoters of previously identified Ume6 targets (top) and in the promoters of the 5’-extended transcripts identified in this study (bottom). Representative images from one of three replicates are shown. **G-H.** For genes with both Ume6 enrichment and a URS1 motif +/- 300 bp from their TSS, the degree of conservation across the 100 bp +/- the URS1 motif center was determined by phastcons within the *sensu stricto* genus. **G.** Metagene analysis for degree of conservation in the promoters of previously identified Ume6 targets compared to the promoters of the 5’-extended transcripts identified in this study. **H.** Heatmap of degree of conservation in the promoters of previously identified Ume6 targets (top) and in the promoters of the 5’-extended transcripts identified in this study (bottom).

The LUTI-based mechanism is neither limited to *NDC80*, nor restricted to budding yeast meiosis. After the discovery of *NDC80^LUTI^*, other LUTIs have been identified in budding yeast and human cells (Cheng et al., 2018; Van Dalfsen et al., 2018; Hollerer et al., 2019). Genome-wide analysis has revealed at least 380 LUTIs that are expressed at various meiotic stages in budding yeast (Cheng et al., 2018). For these loci, transcript levels inversely correlate with protein expression (Cheng et al., 2018). Furthermore, during the endoplasmic reticulum unfolded protein response, the conserved transcription factor Hac1 induces at least 15 LUTIs, most of which are required for downregulating genes involved in cellular respiration (Van Dalfsen et al., 2018). Finally, the *MDM2* oncogene is regulated by a LUTI-based mechanism in human cells (Hollerer et al., 2019). The prevalence of LUTIs in budding yeast and their discovery in humans, along with the pervasive use of alternative transcription start sites (TSSs) and the frequency of uORFs in many organisms all point to the conservation of LUTI-based gene repression across vast evolutionary time (Batut et al., 2013; Calvo et al., 2009; Chew et al., 2016; Johnstone et al., 2016; Kimura et al., 2006).

Previous studies of LUTIs either provided a deep investigation of a single gene or involved analysis of genome-wide datasets to functionally identify LUTIs by looking for instances in which mRNA and protein levels inversely correlate. While both approaches have been successful, many questions remain to be answered: First, why do only some 5’-extended transcripts appear to repress gene expression while others do not? Is the difference due to lack of translational repression, transcriptional repression or both? Second, when gene repression does occur, does it involve a common mechanism? Lastly, what are the key features of the LUTI-based mechanism on a global scale?

In this study, we developed a pipeline to identify and functionally characterize LUTIs in budding yeast meiosis. Using transcript leader sequencing (TL-seq) in conjunction with RNA sequencing on Nanopore arrays (direct RNA-seq), we discovered 74, 5’-extended transcript isoforms as candidate LUTIs. These transcripts are expressed specifically during meiotic prophase but not in premeiotic conditions. 80% of these transcripts are directly regulated by the Ime1-Ume6 transcription factor complex with consensus binding motifs conserved across the budding yeast *sensu stricto* genus. Importantly, uORFs are present in 97% of the 5’-extended transcripts and that almost all of them are bona fide LUTIs based on uORF-dependent translational repression. Transcriptionally, the outlook is more complex: LUTI transcription leads to downregulation of the ORF-proximal transcript in only 50% of cases. Chromatin modifications, specifically histone 3 lysine 36 trimethylation (H3K36me3), and changes in nucleosome position are among the strongest predictors of LUTI-based repression. Finally, higher LUTI expression leads to a greater repression of the proximal promoter-derived transcripts. We conclude that translational repression, chromatin state changes, and degree of LUTI expression act in a combinatorial manner to determine the extent of LUTI-based gene repression. Our findings provide a roadmap to uncover similar types of gene regulation in many other contexts.

## RESULTS

### The combined use of transcript leader and Nanopore sequencing identifies 74 potential LUTIs in meiotic prophase

In order to determine the prominent features of LUTI-based repression, we first developed a pipeline to identify meiotic mRNAs with 5’ extensions (Figure 1B). For cell synchronization, we adopted a previously established protocol in which the expression of two early meiotic regulators, Ime1 and Ime4, are controlled by a copper inducible *CUP1* promoter (Berchowitz et al., 2013). Using this system, we aimed to uncover all 5’-extended mRNAs that are expressed in meiotic prophase (2 hours after induction of *IME1* and *IME4*), but not in premeiotic phase (no induction of *IME1* and *IME4*). Identification of the 5’-extended isoforms relied on data from two orthogonal sequencing techniques. The first method, TL-seq, involves selectively sequencing the 5’ end of a transcript (Arribere and Gilbert, 2013; Malabat et al., 2015). TL-seq allowed for the identification of transcription start sites (TSSs) that robustly increased in expression as cells transitioned from premeiotic phase to meiotic prophase (Figure 1B, 1C). The second technique, Nanopore sequencing, can directly sequence an entire RNA transcript as a single read (Garalde et al., 2018). Nanopore sequencing confirmed the instances in which TSSs identified by TL-seq produced transcripts that elongate across an entire open reading frame (ORF) rather than being terminated early (Figure 1-supplement 1). Lastly, to distinguish LUTIs from the canonical, meiosis-specific mRNAs, we restricted our calls to loci in which the TSS identified by TL-seq was upstream of a second TSS located proximal to the corresponding ORF (Figure 1B, 1C). These analyses resulted in the identification of 74 potential LUTIs, 28 antisense transcripts, 65 intergenic transcripts and 74 intragenic transcripts that were specifically induced in meiotic prophase (Supplemental Table 1 and 2).

### The vast majority of the putative LUTIs are regulated by the same meiotic transcription factor

With a list of 74 LUTI candidates, we sought to determine those that were regulated by a common transcription factor. A search for enriched regulatory motifs in the promoters of 5’-extended transcripts produced a single significant hit matching the URS1 consensus motif (Figure 1D, Supplemental Table 2, 50/74 sequences, combined match p-value < 0.05) (Sumrada and Cooper, 1987). In mitosis, this motif is bound by Ume6, which functions as a transcriptional repressor (Park et al., 1992). However, in meiosis, Ume6 becomes a transcriptional co-activator upon binding to Ime1, culminating in the expression of genes necessary for meiotic entry (Bowdish et al., 1995). The Ime1-Ume6 complex is known to regulate the expression of *NDC80^LUTI^* (Chen et al., 2017). Additionally, Ume6 has been shown to repress 5’-extended transcripts during mitosis at the *BOI1, CFT2,* and *RTT10* loci (Lardenois et al., 2015; Liu et al., 2015). Due to the presence of a URS1 motif upstream of these 5’-extended mRNAs, we hypothesized that Ime1-Ume6 may play a regulatory role at many of the loci producing 5’ extensions during meiotic prophase.

To investigate how many of the URS1 sites are in fact bound by Ume6, we performed chromatin immunoprecipitation followed by deep sequencing (ChIP-seq). Enrichment of Ume6 was present at 61 of the 74 candidate LUTI promoters (q-value < 0.001, fold enrichment > 4). The rate of enrichment was similar in this set of genes compared to a set of previously identified Ume6 targets (canonical, Figure 1E-H) (Williams et al., 2002). Both of these groups were far more enriched with Ume6 than genes not in these lists (Figure 1E and 1F, Figure 1-supplement 2, Figure 1-supplement 3).

We further examined URS1 motif conservation in putative LUTI promoters as a means to assess functional significance. Using an alignment of 5 yeast species in the *sensu stricto* clade (*S. cerevisiae*, *S. paradoxus*, *S. mikatae*, *S. kudriavzevii*, and *S. bayanus*), conservation of the region +/- 100 bp around the URS1 motif was calculated for all sites that had both a URS1 motif and were bound by Ume6. For the previously defined canonical Ume6 targets and the regions around the newly identified 5’-extensions, conservation sharply increased around the URS1 motif (Figure 1G and 1H); providing strong evidence that 33 of the identified 5’-extended transcript isoforms have strongly conserved URS1 binding sites, and their regulation by Ume6-Ime1 is likely functional (Figure 1-supplement 2).

### Transcript isoform diversity and the translational regulation of 5’-extended isoforms

LUTIs were originally identified based on an inverse correlation between transcript and protein levels (Chen et al., 2017; Cheng et al., 2018). Of the 380 LUTIs identified throughout meiosis by Cheng et al. (2018), 32 (8.4%) of them were found to be expressed during the meiotic prophase time point collected in this study. The remaining 42 of our 74 candidate LUTIs were previously unidentified. While ten of the false-negatives were missed in part due to lack of a quantifiable protein measurement, the remainder were likely overlooked due to differences in the method of identification (Cheng et al., 2018). Our pipeline detects all 5’-extended transcripts, regardless of their correlation with protein abundance. Such newly identified LUTI-candidates are potentially very interesting, because they might reveal new biological insight into how frequently candidate LUTIs repress gene expression and under what circumstances this repression occurs.

It is difficult to answer these questions using traditional RNA-seq, because the transcript produced from the ORF-proximal promoter, hereon referred to as the PROX isoform, does not have any unique identifying sequence compared to the transcript produced from the ORF-distal promoter. By sequencing the most 5’ end of a transcript with TL-seq, this was no longer a limitation. We performed TL-seq and RNA-seq in parallel on samples collected during premeiotic phase and meiotic prophase. RNA-seq and TL-seq measurements (in which the closest TSS to ORF was quantified) correlated well for most genes, demonstrating that TL-seq can quantitatively estimate transcript abundances (Spearman’s rank correlation coefficient, ρ= 0.771 [premeiotic], ρ= 0.732 [meiotic prophase], Figure 2A and 2B, teal). However, for genes with 5’-extended transcripts, this correlation was poor in meiotic prophase (ρ= 0.419, Figure 2B, orange). Even more strikingly, when the fold-change of expression between meiotic prophase and premeiotic stages was taken into account, there was no correlation between TL-seq and RNA-seq for genes with 5’-extended transcripts (ρ= 0.04, Figure 2C, orange). This suggests that TL-seq is a more robust method than RNA-seq to specifically quantify the 5’-extended and PROX isoforms.

**Figure 2.**
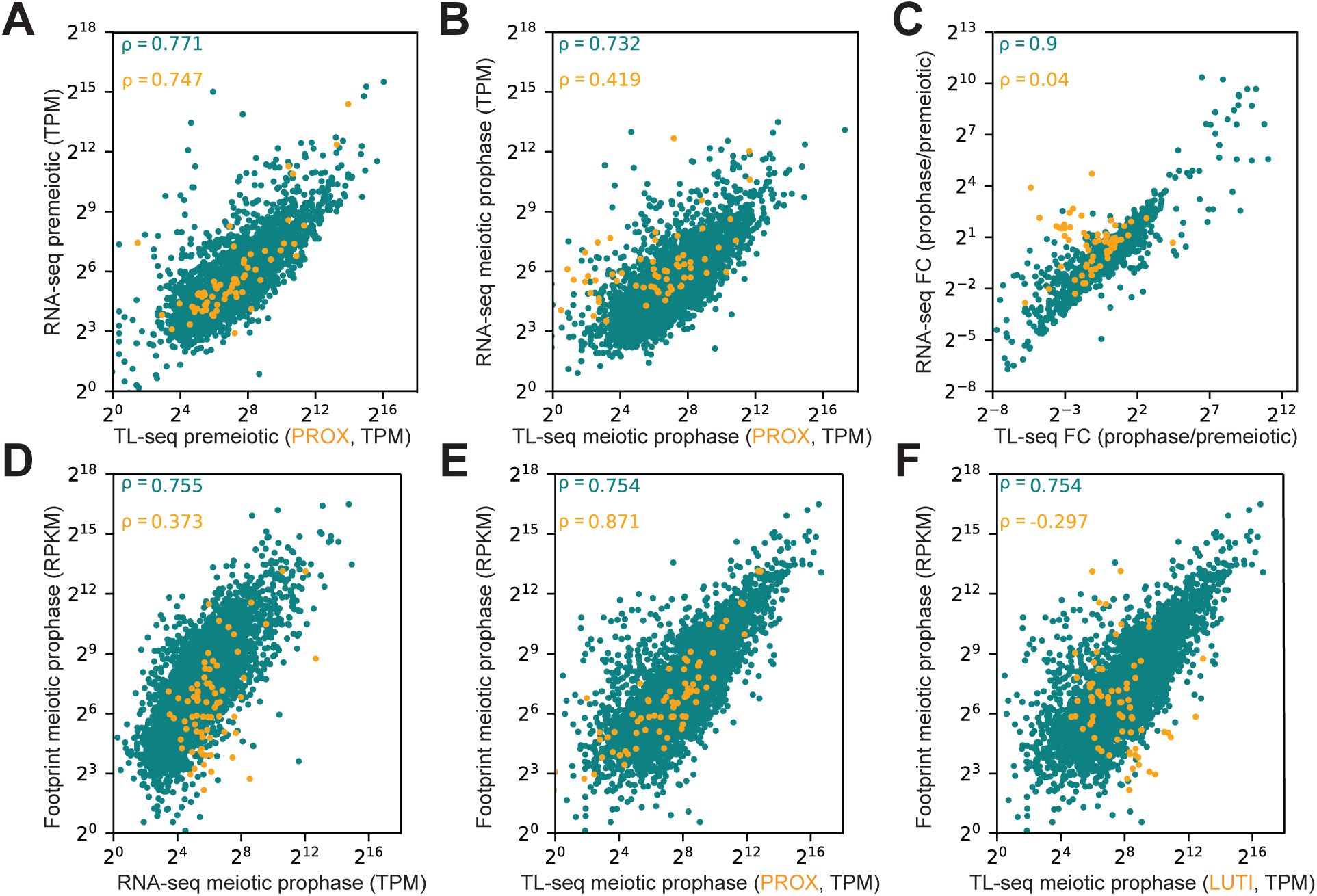
Relationship between transcript isoform diversity and translational regulation of 5’-extended isoforms. **A-B.** Scatterplot of RNA-seq and TL-seq for genes with (orange) and without (teal) 5’-extensions. For genes with 5’-extensions, only the PROX transcript is quantified. Cells were induced to undergo meiosis and collected during meiotic prophase. Experiments were performed in duplicate. The TL-seq and RNA-seq were taken from matched RNA samples. Transcripts per million were quantified by salmon for RNA-seq and cageR for TL-seq. Note that the matched TL-seq data was not sequenced to the same depth as the original TL-seq data used for isoform discovery, so 11 loci with 5’-extensions were not quantified. Spearman’s rank correlation coefficients (ρ) are displayed in the upper left corner for A-F. Cells with the *pCUP1-IME1/pCUP1-IME4* meiotic induction system (*UB14584*) were collected for TL-seq and RNA-seq after (**A**) 2 hours in sporulation medium (SPO), before induction of meiosis, or (**B**) after 4 hours in sporulation medium (SPO), which corresponds to 2 hours after induction of meiosis by 50 µM CuSO_4_. **C.** The fold-change (FC) by which gene expression changes as the cells enter meiotic prophase from premeiotic stage is quantified by DESeq2. Spearman’s rank correlations were calculated for genes with (orange) and without (teal) 5’-extensions. For genes with 5’-extensions, only the PROX transcript is quantified. **D-F.** Scatterplot of translation as measured by ribosome footprints vs (**D**) RNA-seq (in duplicate) and (**E-F**) TL-seq (in triplicate, the TL-seq replicates used to initially identify loci with 5’-extensions) for all genes without 5’-extensions (teal) and genes with 5’-extensions (orange). In (**E**), only the PROX transcript is quantified for genes with 5’-extensions, and in (**F**) only the LUTI transcript is quantified for genes with 5’-extensiosn. The ribosome profiling data come from the 3h time point in Cheng et al. 2018. Cells with the *pCUP1-IME1/pCUP1-IME4* meiotic induction system (*UB14584*) were collected for TL-seq and RNA-seq after 4 hours in sporulation medium (SPO), which corresponds to 2 hours after induction of meiosis by 50 µM CuSO_4_.

We next set out to compare the correlation between ORF translation and transcript abundance, with the added ability to distinguish between the two transcript isoforms produced in meiotic prophase. We took advantage of a published matched ribosome profiling and RNA-seq dataset (Cheng et al., 2018). The meiotic prophase RNA-seq dataset from this study best matched the 3-hour time point from Cheng et al; therefore we used the ribosome profiling data from the 3-hour time point for all further analyses (Figure 2-supplement1). For genes without 5’-extensions, the distribution between RNA-seq and ribosome footprints looked very similar to the distribution between TL-seq and footprints (ρ= 0.755 vs ρ= 0.754, Figure 2D-F, teal). However, for the subset of genes with 5’ extensions, there were fewer footprints per unit mRNA relative to the rest of the transcriptome, indicating that these transcripts are poorly translated (ρ= 0.373, Figure 2D, orange). Remarkably, the PROX transcripts quantified by TL-seq were highly proportional to translation measurements (ρ= 0.871, Figure 2E, orange), while the 5’-extended transcripts showed negative correlation (ρ= −0.297, Figure 2F, orange). This analysis demonstrates that transcript isoform-specific measurements are far more accurate in assessing translational status than RNA-seq. Furthermore, it provides additional evidence that the identified 5’-extended transcripts, as a whole, do not productively translate the ORFs contained within them.

### Prevalence of uORFs in meiotic LUTIs

We next investigated how the candidate LUTIs were translationally impaired. While 5’-leader mediated translational repression can occur through multiple ways including secondary RNA structure, RNA modification, protein-RNA interactions (reviewed in Hinnebusch et al., 2016), based on the abundance of uORFs in meiosis (Brar et al., 2012) as well as the uORF-dependent translational repression of *NDC80^LUTI^*(Chen et al., 2017), we hypothesized that uORFs would play a dominant role in dampening ORF translation from other LUTIs. The number of ATG codons in the region between the distal TSS and the PROX TSS was used to determine uORF abundance in candidate LUTI mRNAs. Only ATG uORF start codons were counted, because current evidence has not determined whether uORFs with near-cognate start sites prevent translation of the downstream ORF (Brar et al., 2012; Guenther et al., 2018).

Our analysis revealed that 72 of the 74 candidate LUTIs had at least 1 ATG uORF and that 64 of these 72 had between 4 and 17 ATGs (Figure 3A). The two genes without an ATG uORF (*PLM2* and *YCL057C-A*), displayed an increase in TE (translational efficiency= ribosome footprints RPKM / transcript abundance RPKM) upon entry into meiosis, indicating that these genes should not be categorized as LUTIs (Figure 3-supplement 1). The observation that the 5’-extended transcripts without any ATG uORFs were better translated further strengthens the argument that ATG uORFs are causal for translational repression.

**Figure 3.**
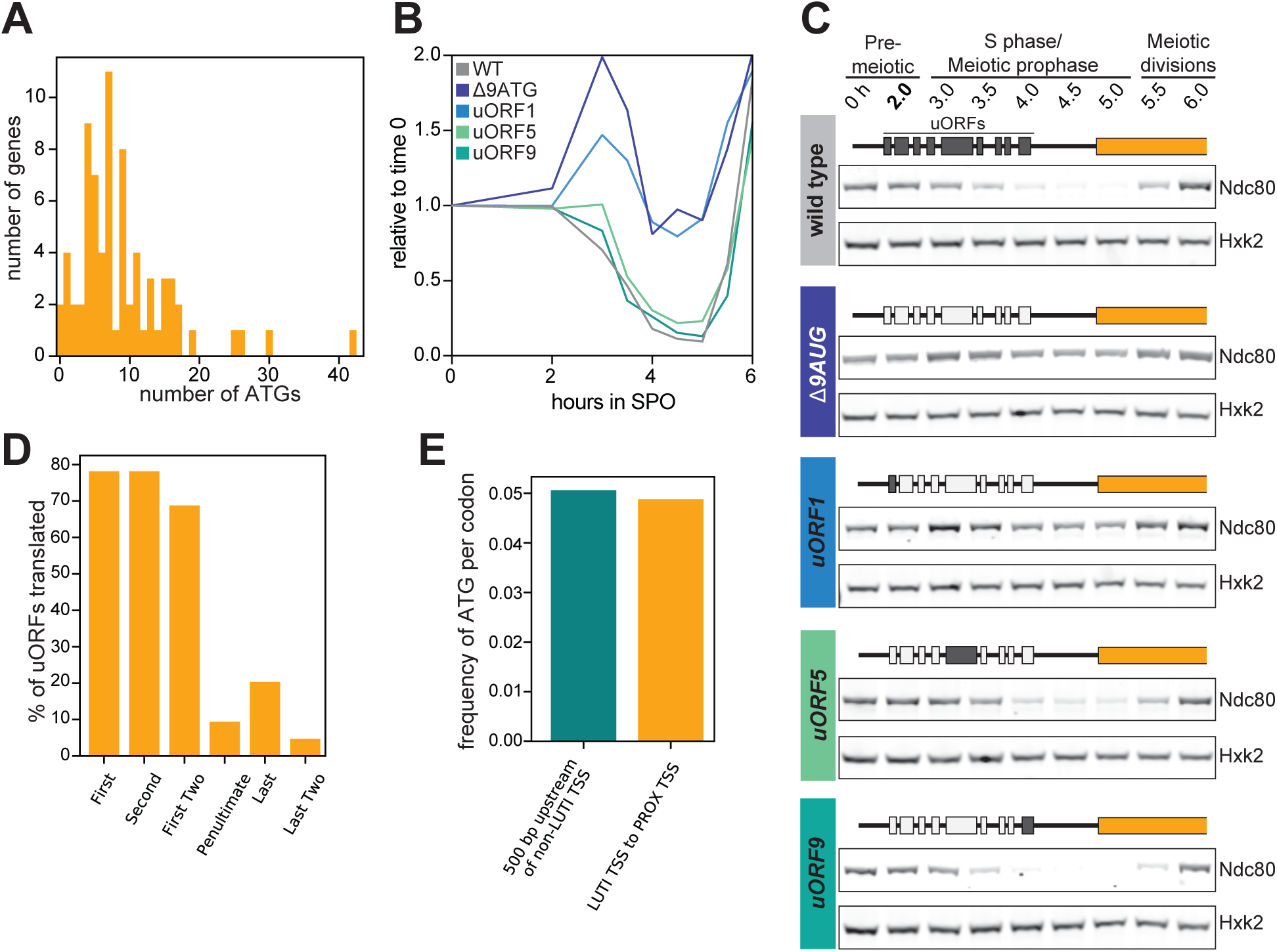
uORF-based translational repression is prevalent in LUTIs. **A.** A histogram of the number of ATGs found in the region between the proximal and the distal TSSs at loci with LUTI mRNAs. **B-C.** A single uORF can be sufficient to repress translation. Cells with the *pCUP1-IME1/pCUP1-IME4* meiotic induction system and 3V5-tagged *NDC80* were induced to undergo meiosis by the addition of 50 µM CuSO_4_ after 2 hours in SPO. Strains used included wild type (*UB6190*), a strain in which all of the ATGs in the *NDC80^LUTI^* leader were mutated to ATC (*Δ9AUG: UB6183*), and strains in which only the first (*uORF1: UB10579*), fifth (*uORF5: UB10581*), or ninth (*uORF9: UB10583*) ATG was left intact. Immunoblots were performed on samples collected between 0-6 hours in SPO. Immunoblots were performed with a α-V5 antibody to recognize the 3V5-tagged Ndc80. Hxk2 was used as a loading control. The blots represent one of two replicates. **B.** Quantification of the western blots. Signal at each time point was first normalized to the Hxk2 loading control and then to the first time point (time 0). The quantification represents one of two replicates. **C.** Immunoblots from quantification in (**B**). In the illustrations above each blot, the darkened uORFs represent those that have an ATG in the respective strain. **D.** The frequency of uORF translation. In LUTIs containing at least 4 uORFs, the translation state of the two most 5’ and the two most 3’ AUG uORFs were assessed. If > 4 footprints were observed in the first 6 codons, excluding the AUG start codon, a uORF was considered translated. In cases where two uORFs were evaluated, the set was considered translated if both uORFs independently had > 4 footprints. **E.** The frequency of ATGs in the region between the LUTI TSS and the ORF-proximal TSS was compared to the region 500 bp upstream of TSS that do not express LUTIs in meiotic prophase.

In the cases when an extended transcript contained a single uORF, the effect on translational efficiency was variable. The TE changed very little for 2 genes (*ELO1* and *COX16*), possibly because the LUTI was the minor isoform. The LUTI was the major isoform in the other 2 cases, and the TE decreased dramatically by ∼200-fold for *MNE1* and ∼20-fold for *ULP1* (Figure 3-supplement 1), suggesting that a single uORF was sufficient to inhibit translation initiation at downstream AUGs in these examples. Based on these analyses, all of the 5’-extended transcripts containing at least a single ATG uORF will be referred to as LUTIs going forward.

We directly tested the ability of a single uORF to repress translation of the downstream ORF at the *NDC80* locus. *NDC80^LUTI^* normally contains 9 uORFs in its 5’ leader. When all 9 ATGs were mutated to ATCs (*Δ9AUG*), Ndc80 protein was translated from the LUTI (Figure 3B and 3C), consistent with previous work (Chen et al., 2017). Strains were constructed in which the ATG of either the first, fifth, or ninth uORF was the sole ATG-initiated uORF in the 5’ leader. Having the first uORF alone resulted in similar Ndc80 protein levels to what was observed in the *Δ9AUG* strain; however, LUTIs carrying only the fifth or the ninth uORF repressed Ndc80 translation just as efficiently as the wild type *NDC80^LUTI^* (Figure 3B and 3C). We conclude that a single uORF can be sufficient to cause translational repression.

Because the presence of uORFs does not always lead to repression, we set out to establish another metric to determine whether LUTIs are translationally repressed. As the uORF number in a 5’-leader increases, the likelihood of repression at the downstream ORF also increases (Calvo et al., 2009; Chew et al., 2016; Johnstone et al., 2016). We predicted that if translational repression occurs, there would be more translation over the most 5’ uORFs and less translation over the uORFs closest to the annotated gene’s coding region. Because uORFs are short and frequently overlapping, it can be difficult to accurately quantify their translation. Instead we determined a threshold of at least 4 footprint counts within the first 6 codons of a uORF to call it as translated. With this metric, almost 80% of the first 2 uORFs in transcripts with at least 4 uORFs were classified as translated (Figure 3D). Compellingly, less than 15% of the last 2 uORFs in those same transcripts were determined to be translated (Figure 3D). Thus, the ribosomes frequently get caught up before scanning across all uORFs. This is consistent with the observation that ATG frequency was not higher in the 5’-extensions compared to the 500 bp upstream of genes not expressing LUTIs in meiotic prophase (Figure 3E). If indeed LUTIs do play important and functional roles in mediating meiotic gene expression, the lack of uORF selection would indicate that the natural frequency of ATGs in intergenic regions is sufficient to result in the necessary degree of translational inhibition. We conclude that uORFs are found in abundance in the 5’-leaders of most LUTIs, and furthermore, they efficiently prevent the ribosome from translating the downstream ORF.

### Variable transcriptional repression by LUTIs

In addition to the translational impairment of LUTIs, transcription of the previously identified LUTIs prevents transcription from the PROX promoters (Chen et al., 2017; Hollerer et al., 2019). To assess the degree of transcriptional repression in the set of LUTIs identified in this study, the abundance of LUTI and PROX transcripts were measured by TL-seq. LUTI levels did not correlate with PROX transcript abundance prior to meiotic entry (ρ= −0.19, Figure 4A); however, a significant negative relationship developed in meiotic prophase (ρ= −0.359, p-value = 1.98 x 10^-3^), associating LUTI expression with a decrease in PROX isoform level (Figure 4B). In addition, the PROX transcript abundance in meiotic prophase was less than in premeiotic stage for a large number of genes, further supporting their transcriptional repression by LUTIs (Figure 4C).

**Figure 4.**
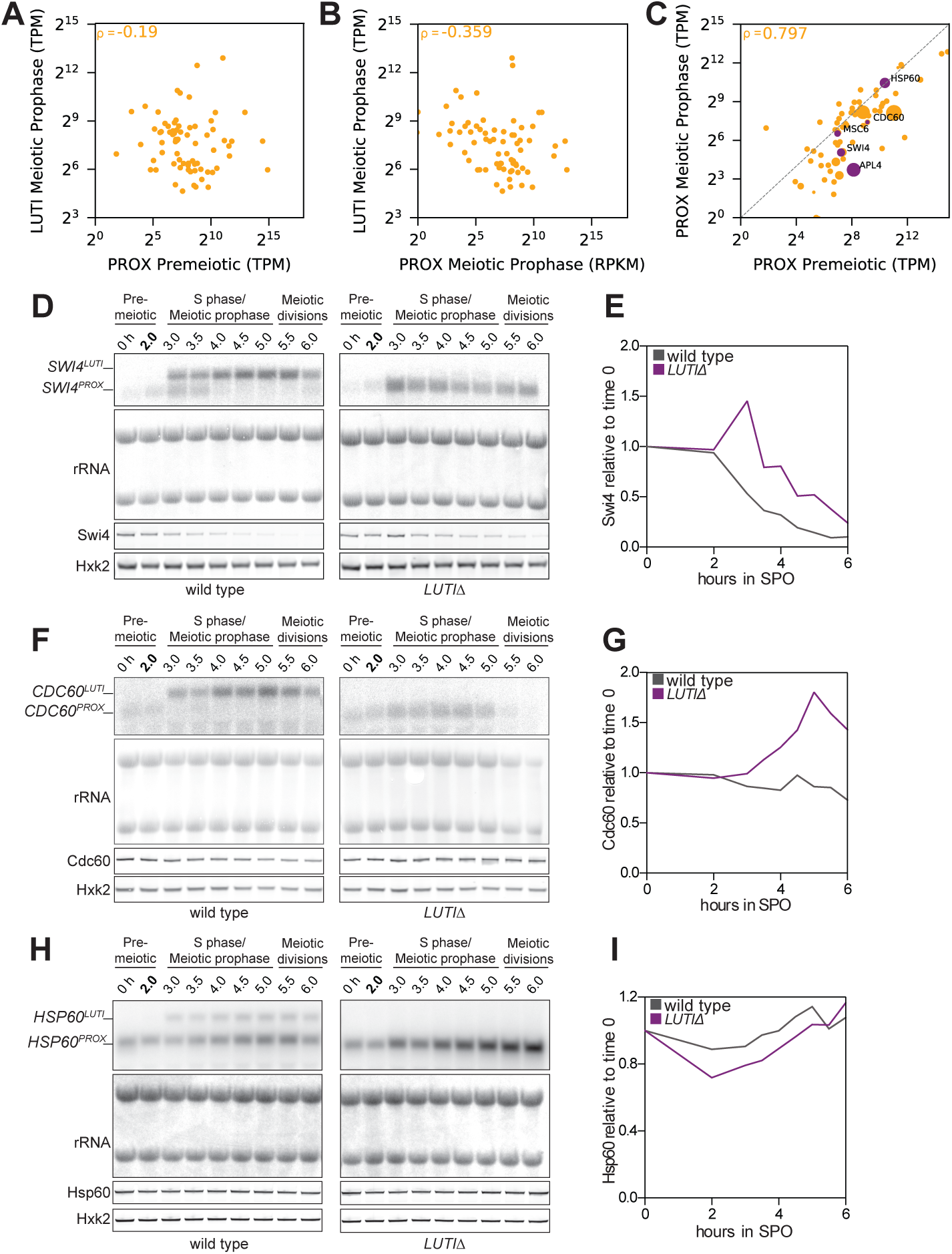
Transcriptional repression by LUTIs. **A.** Scatterplot of LUTI abundance in cells from meiotic prophase vs PROX abundance in cells from premeiotic stage from the *pCUP1-IME1/pCUP1-IME4* meiotic induction system (*UB14584*). The premeiotic time point was taken after 2 hours in SPO, and the meiotic prophase time point was taken after 4 hours in SPO, which corresponds to 2 hours after 50 µM CuSO_4_ addition. The values plotted are from the mean of 3 replicates and are displayed in transcripts per million (TPM) as determined by cageR. The Spearman’s correlation coefficient (ρ) is displayed in the upper left corner. **B.** The same as A except that the PROX transcript abundance was calculated from the meiotic prophase time point. **C.** Visualization of the change in PROX transcript abundance after cells enter meiosis. Scatterplot of PROX abundance in meiotic prophase compared to premeiotic as transcripts per million (TPM). The size of each point correlates to the LUTI abundance in meiotic prophase. The Spearman’s correlation coefficient (ρ) is displayed in the upper left corner. Purple points indicate genes of interest selected for in-depth analysis. **D-I.** LUTI deletion analysis of single genes. Cells with the *pCUP1-IME1/pCUP1-IME4* meiotic induction system were induced to enter meiosis using 50 µM CuSO_4_ and collected after 2 hours. (**D-E**) *SWI4-3V5* tagged strains with either wild type (*UB18175*) or LUTIΔ (*UB18176*) (**F-G**) *CDC60-3V5* tagged strains with either wild type (*UB18185*) or LUTIΔ (*UB18186*), (**H-I**) *HSP60-3V5* tagged strains with either wild type (*UB18336*) or LUTIΔ (*UB18188*). (**D**), (**F**) and (**H**) RNA blots and Immunoblots were performed on samples collected between 0-6 hours in SPO. RNA blots were performed with a probe specific for 3V5 and its linker. Methylene blue detection of rRNA bands was used as a loading control. Immunoblots were performed with a α-V5 antibody to recognize 3V5-tagged proteins. Hxk2 was used as a loading control. These blots represent one of two replicates for *CDC60* and *SWI4*, and a single replicate for *HSP60*. (**E**), (**G**) and (**I**) Quantification of the Immunoblots shown in (**D**), (**F**) and (**H**). 3V5 signal was quantified relative to Hxk2 and then relative to 0 h (time 0). Plots represent one of two replicates for *CDC60* and *SWI4*, and a single replicate for *HSP60*.

To parse out how causative the relationship was between LUTI and PROX expression, five genes were selected for in-depth analysis. Two of them (*SWI4* and *APL4*) had very strongly repressed PROX transcripts in meiotic prophase, two (*MSC6* and *HSP60*) had PROX transcripts present at similar levels in both time points, and one (*CDC60*) had an intermediate amount of PROX repression (Figure 4C). For each gene, transgenes carrying a 3V5 epitope tag with either a normal LUTI promoter (wild type) or a deletion (*ΔLUTI*) were constructed. Cells were synchronized in meiosis to track transcript isoforms and protein abundance by RNA blotting and immunoblotting, respectively. In all instances, including those loci in which PROX expression holds steady before and after wild type LUTI induction, the LUTI deletion led to an increase in PROX transcript abundance (Figure 4D-I, Figure 4-supplement 1A and 1B). This translated to an increase in protein levels for Swi4 and Cdc60 (Figure 4E and 4G), but not for Apl4, Msc6 and Hsp60, likely due to differential protein stability (Figure 4I, Figure 4-supplement 1A and 1B). This suggests that even when the LUTI-based repression is relieved, additional layers of control, such as posttranslational regulation, can buffer the final protein output.

### The role of chromatin in LUTI-based transcriptional repression

With the knowledge that LUTI expression can play a role in repressing transcription from downstream TSSs at some loci, but not others, we set out to understand what differentiates between the two cases. In both yeast and human cells, LUTI expression leads to an increase in histone 3 lysine 36 trimethylation (H3K36me3) over the proximal gene promoter (Chia et al., 2017; Hollerer et al., 2019). In yeast, *NDC80^LUTI^* expression also results in histone 3 lysine 4 dimethylation (H3K4me2) enrichment (Chia et al., 2017). Both marks are necessary for LUTI-based repression of *NDC80* (Chia et al., 2017). H3K4me2 and H3K36me3 are found at sites of active transcription, with H3K4me2 present in more abundance towards the 5’ end of a transcribed region (Kirmizis et al., 2007; Liu et al., 2005; Pokholok et al., 2005). Despite localizing to regions undergoing active transcription, these marks are not necessarily associated with gene activation. Particularly in budding yeast, H3K36me3 plays a role in repressing transcription initiation within gene bodies (Carrozza et al., 2005; Keogh et al., 2005; Li et al., 2007). The role of H3K4me2 is less well understood, but it is thought to play a role in the induction kinetics of genes at loci with overlapping transcription (Kim and Buratowski, 2009; Kim et al., 2012; Pijnappel et al., 2001).

To determine the change in H3K36me3 and H3K4me2 patterns at LUTI-associated loci, we performed ChIP-seq. In premeiotic cells, the LUTI-containing genes displayed dis-enrichment of both modifications just upstream of the PROX TSS (Figure 5A and 5B, top panel, Figure 5-supplement 1, Figure 5-supplement 2). Interestingly, H3K36me3 was enriched in meiotic prophase specifically over the promoters of those genes expressing LUTIs (Figure 5A, bottom panel). Strikingly, those genes whose PROX transcript is most repressed (fold change < 0.25, n=21) have the highest levels of H3K36me3 over their proximal TSS (Figure 5A, bottom panel, Supplemental Table 3). In contrast, genes that have associated LUTIs, but do not experience a decrease in the abundance of the PROX transcript (log2 fold change > 0, n=18), have only a minor increase in H3K36me3 levels (Figure 5A, bottom panel, Supplemental Table 3). For H3K4me2, a moderate increase in the chromatin modification is observed over the PROX promoters of only those genes that are most repressed in meiotic prophase (Figure 5B, bottom panel, Supplemental Table 3). Thus, while increased H3K36me3 is a strong predictor for LUTI-based repression, H3K4me2 appears to be altered in a more limited number of instances, such as in the case for *NDC80*.

**Figure 5.**
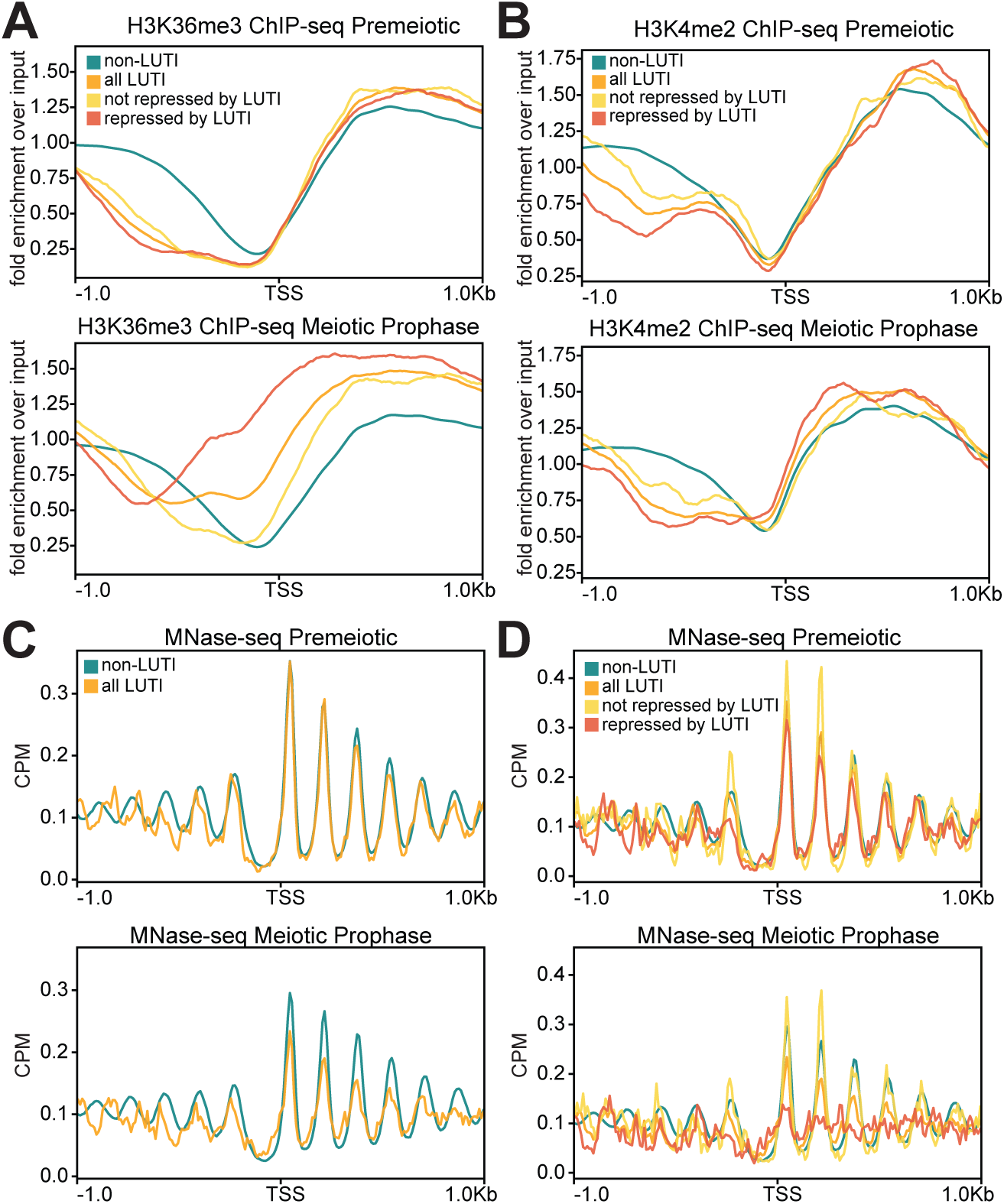
The role of chromatin modifications and nucleosome position in LUTI-based transcriptional repression. **A.** Metagene analysis of H3K36me3 ChIP-seq in premeiotic and meiotic prophase stages. The ChIP was performed in cells with the *pCUP1-IME1/pCUP1-IME4* meiotic induction system *(UB14584*). The premeiotic time point was taken after 2 hours in SPO, and the early meiotic time point was taken after 4 hours in SPO, which corresponds to 2 hours after 50 µM CuSO_4_ addition. The repressed LUTI genes include the subset of genes with a fold-change in PROX abundance of < 0.25 between the early meiotic and premeiotic time points (n=21). The non-repressed LUTI genes are those with a fold-change in PROX abundance of > 1 (n=18). The plot is centered around the PROX TSS. The images are from one of three replicates. **B.** Same as (**A**) but for H3K4me2. The images are from one of three replicates. **C.** Metagene analysis of MNase-seq in premeiotic and meiotic prophase cells. Performed with the same strain and time points as in (**A**). The images are from one of three replicates. **D.** Same as (**C**), but including the non-repressed and repressed groups of genes from (**A**). The images are from one of three replicates.

In addition to methylation states of the histones around the PROX TSS, the presence of nucleosomes can occlude the binding of transcription factors and other machinery required for transcription initiation (Klemm et al., 2019; Venkatesh and Workman, 2015). Evidence from the case of *NDC80^LUTI^* indicates that upstream transcription leads to nucleosome repositioning around the ORF-proximal promoter thereby shrinking the nucleosome-depleted region (Chia et al., 2017). By performing micrococcal nuclease digestion followed by deep sequencing (MNase-seq), we tracked genome-wide changes to nucleosome positions. In meiotic prophase, but not in premeiotic stage, the nucleosome peaks decreased while signal in the valleys increased specifically for the LUTI-associated genes, thus increasing the “fuzziness” of nucleosomes (Figure 5C) (Chen et al., 2013). The effect was strongest for loci with the greatest degree of PROX transcript repression (Figure 5D). In that subset of 21 genes, the nucleosome position was so disrupted that a consensus nucleosome periodicity could not be identified (Figure 5D, bottom panel). Examining such a small subset of genes might make it hard to observe strong periodicity; however, in a similarly small subset of genes (n=18), loci with non-repressive LUTIs, periodic nucleosome positioning was still present (Figure 5D, bottom panel). Therefore, the complete lack of periodicity at loci with repressive LUTIs, was most likely due to variability in the extent of repositioning at each locus (Figure 5-supplement 3). For this reason, we conclude that robust nucleosome repositioning occurs over the promoters of genes that are most subject to LUTI-based transcriptional repression.

### The features defining LUTI-based transcriptional repression

In addition to H3K36me3 enrichment and increased nucleosome occupancy over proximal promoters, we expected that other variables could play a role in LUTI-based transcriptional repression. Stemming from previous work showing that the distance between promoters with overlapping transcription affects the mechanism of transcriptional repression in cells undergoing carbon source shifts (Kim et al., 2017), we considered the importance of the distance between the PROX and the LUTI TSSs in mediating PROX transcript repression. The LUTI abundance and the length of the gene were also considered, as well as changes to the +1 and –1 nucleosome positions and “fuzziness.” We found that an increase in H3K36me3 (ρ= −0.492, p-value = 1.10 x 10^-5^), high LUTI levels (ρ= −0.456, p-value = 5.60 x 10^-5^), +1 nucleosome peak moving toward the nucleosome-depleted region (ρ= −0.423, p-value = 2.13 x 10^-4^), and an increase in +1 nucleosome “fuzziness” (ρ= −0.341, p-value = 3.41 x 10^-3^) all displayed significant negative correlation with the log2 fold-change of PROX transcript abundance (Figure 6A, Figure 6-supplement 1). However, changes at the –1 nucleosome, H3K4me2 and the distance between the TSSs had no significant correlation (Figure 6A, Figure 6-supplement 1). Altogether, these analyses helped distinguish a set of important factors involved in LUTI-based transcriptional repression.

**Figure 6.**
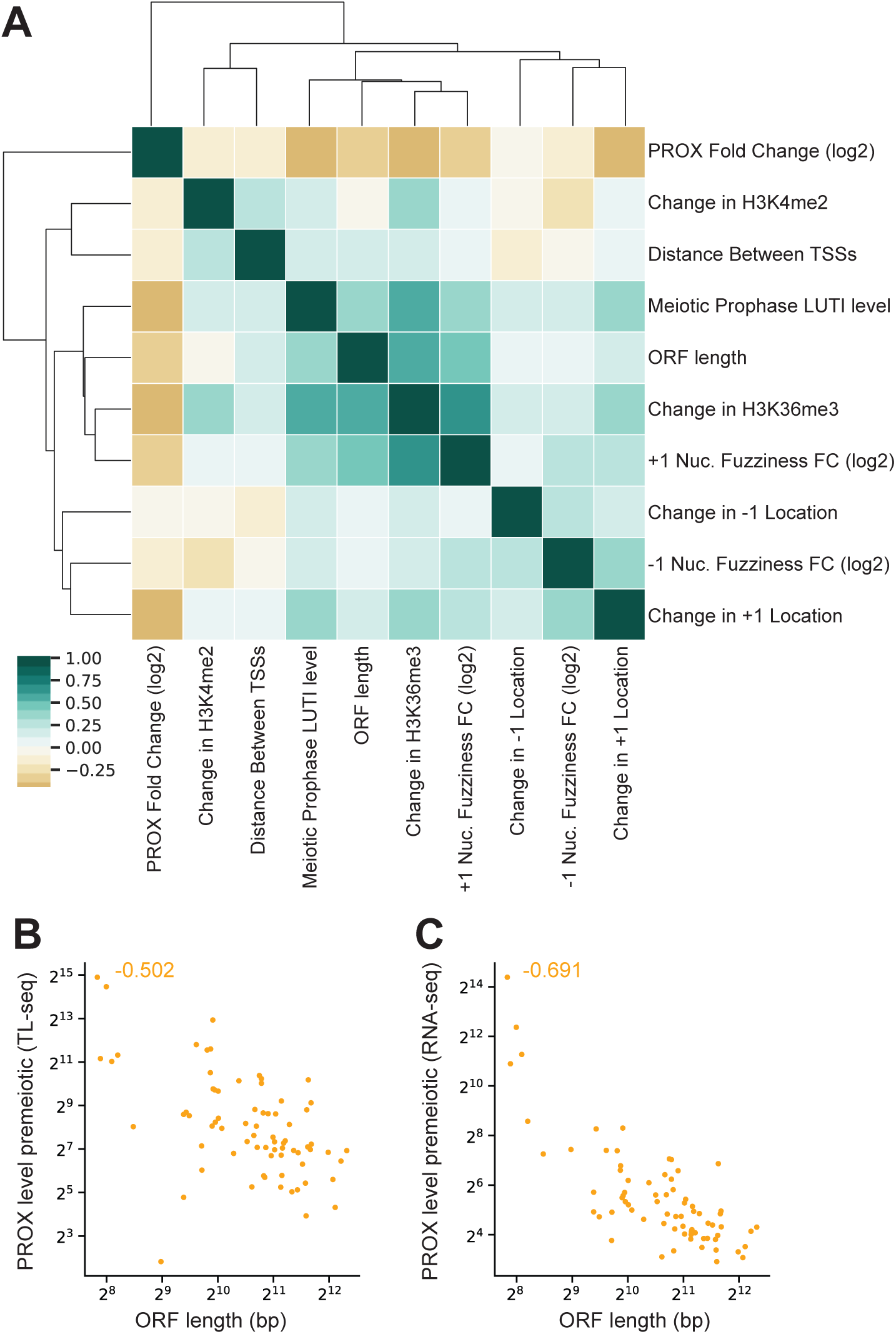
The key features that predict LUTI-based transcriptional repression. **A.** Clustermap of possible features associated with LUTI-based repression. Features were clustered by Spearman’s correlation coefficient (ρ). **B.** Scatter plot of the correlation between ORF length and PROX abundance in premeiotic cells as quantified by TL-seq. The Spearman’s correlation coefficient (ρ) is displayed in the upper right corner. **C.** Same as in (**B**), but the PROX abundance was calculated from RNA-seq data.

### The LUTI and PROX promoter strengths together dictate LUTI-based transcriptional repression

Surprisingly, we also found that the longer a gene’s coding sequence was, the more likely it was to have an abundant LUTI, an increase in H3K36me3, greater +1 nucleosome fuzziness, and an increased likelihood of being repressed by a LUTI (Figure 6A). Coincidentally, we noticed that in the set of genes with meiotic LUTIs, the shorter genes had higher PROX transcript abundances than did longer genes (Figure 6B and 6C). Could it be that the promoters of strongly expressed genes are better able to continue transcribing their gene products even in the presence of LUTI mRNAs? We decided to investigate this possibility further.

If a more robustly transcribed PROX isoform is resistant to repression by LUTIs, it follows that increasing LUTI expression or decreasing PROX expression could shift the balance in favor of LUTI-based repression. To test this hypothesis, we focused on *HSP60*, a LUTI carrying gene whose PROX isoform is highly expressed both in premeiotic stage and in meiotic prophase. We engineered a reporter construct in which the *HSP60* regulatory region (−501 to −1 bp upstream of the *HSP60* ATG) was upstream of a short-lived green fluorescent protein (ubiGFP). ubiGFP has a half-life of ∼ 7 min (Houser et al., 2012), which restricts its detection to the window when the PROX transcript is expressed. To enable titratable expression of LUTI, an inducible LexA/LexO system was used (Ottoz et al., 2014). The endogenous *HSP60^LUTI^* promoter was replaced with 8 copies of the LexO operator fused to a *CYC1* minimal promoter (Figure 7A). A β-estradiol responsive LexA-ER-AD fusion protein could thus control the expression of LUTI. After induction of sporulation, cells were treated with 0, 2 or 5 nM β-estradiol, and samples were subsequently collected at different time points. As the dose of β-estradiol increased, so did the levels of LUTI (Figure 7B). Furthermore, the increased LUTI expression corresponded with a dose-dependent increase in H3K36me3 over the *HSP60^PROX^* promoter (Figure 7C). Importantly, as the expression of LUTI increased, the abundance of ubiGFP simultaneously decreased (Figure 7D and 7E). The dose-dependent decrease in ubiGFP levels was also confirmed during mitotic growth in rich media (Figure 7H and 7I). We conclude that changes to the strength of LUTI expression can lead to a decrease in protein abundance even at a strong PROX promoter.

**Figure 7.**
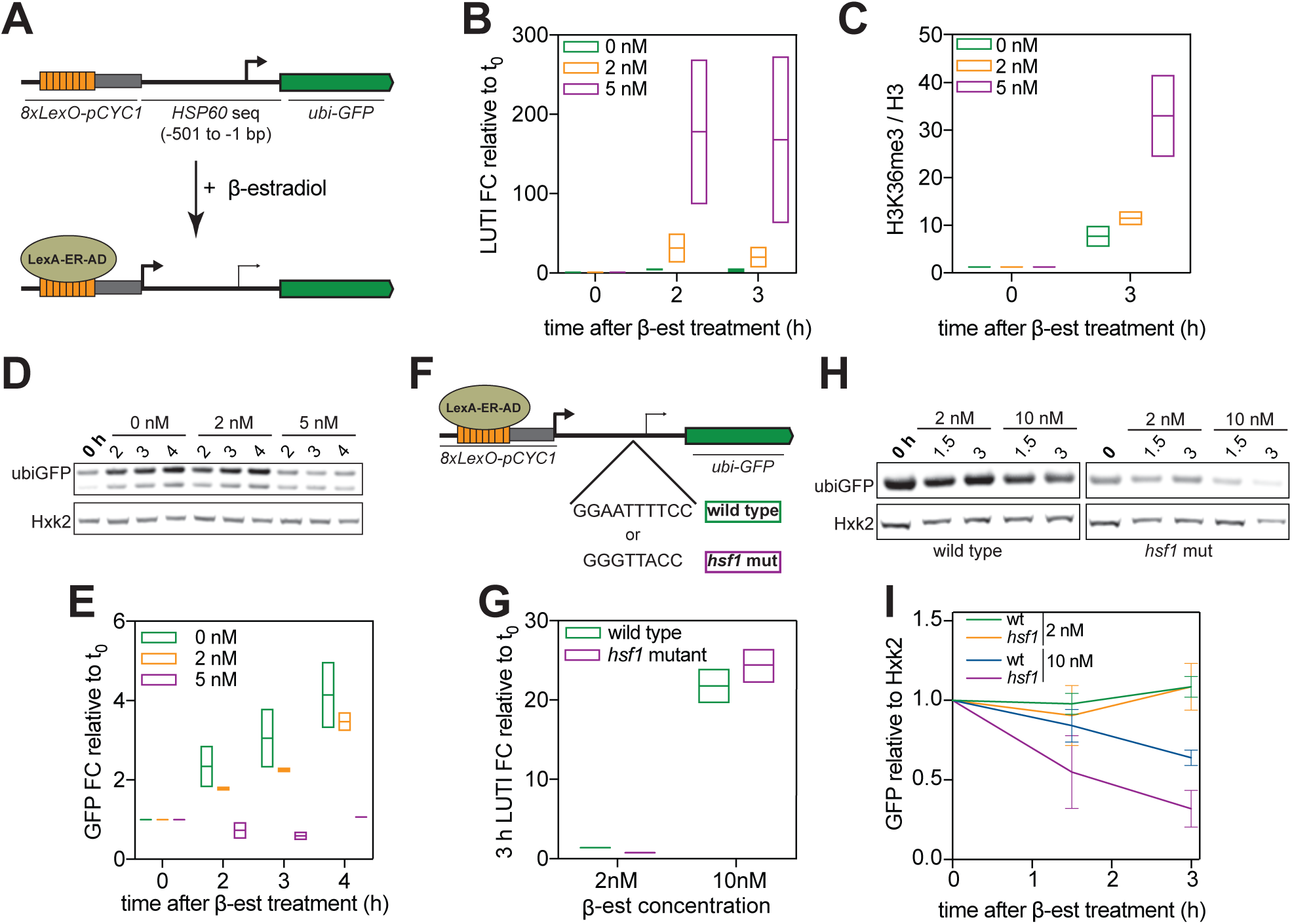
Changes in LUTI or PROX expression both affect LUTI-based repression of a reporter gene. **A.** Illustration of the reporter construct used. The *HSP60^LUTI^* promoter was replaced with 8xLexO and a minimal *CYC1* promoter. This promoter can be induced upon addition of β-estradiol in cells harboring a LexA-ER-AD fusion protein. B112 (219 amino acids), a short unstructured acidic peptide encoded by *Escherichia coli*, is used as an activation domain (AD) for the chimeric transcription factor LexA-ER-AD. A GFP protein with an N-terminal ubiquitin fusion followed by a tyrosine residue (ubi-GFP) is subject to N-end rule degradation following ubiquitin cleavage. This short-lived reporter was engineered into the construct in place of the *HSP60* ORF. **B-E.** Cells with the *pCUP1-IME1/pCUP1-IME4* meiotic induction system in combination with a LexA-ER-AD/8xLexO inducible promoter regulating the *HSP60^LUTI^* upstream of ubi-GFP (*UB19257*) were induced to undergo meiosis with 50 µM of CuSO_4_. *HSP60-ubiGFP^LUTI^* was not induced (0 nM), induced with 2 nM, or induced with 5 nM of β-estradiol after 2 hours in SPO (corresponds to time 0). **B.** RT-qPCR of *HSP60-ubiGFP^LUTI^*. Transcript abundance was quantified using a primer set that spans from the region immediately upstream of the *HSP60* PROX TSS until the beginning of ubiquitin. Quantification was performed in reference to the levels of the meiotic housekeeping gene *PFY1* and then normalized to the 0 h time point. FC=fold change. Experiments were performed in duplicate, and the range is displayed. **C.** H3K36me3 ChIP was performed before induction and 3 hours after induction with β-estradiol, corresponding to 5 hours in SPO. Enrichment was quantified over H3 abundance using the same primer set as in (**B**). Experiments were performed in duplicate, and the range is displayed. **D.** Immunoblot was performed using an α-GFP antibody. Hxk2 was used as a loading control. Image represents one of two replicates. **E.** Immunoblot in (**D**) was quantified relative to Hxk2 and then relative to the time of β-estradiol addition. FC=fold change. **F**. A version of the construct in (**A**) was generated with a mutated Hsf1 binding site (GGAATTTTCC to GGGTTACC). **G-I.** Cells from W303 strain background harboring the *8xLexO-HSP60-ubiGFP* construct with either a wild type (*UB18838*) or *hsf1* (*UB20485*) mutant PROX promoter were collected during exponential mitotic growth before and after induction of *HSP60-ubiGFP^LUTI^.* The induction was performed with either 2 nM or 10 nm of β-estradiol. **G.** RT-qPCR of *HSP60-ubiGFP^LUTI^*. Transcript abundance was quantified using a primer set that spans from the region immediately upstream of the *HSP60* PROX TSS until the beginning of ubiquitin. Quantification was performed in reference to the levels of *ACT1* and then normalized to the time of β-estradiol addition. FC=fold change. Experiments were performed in duplicate, and the range is displayed. **H.** Immunoblot was performed using an α-GFP antibody. Hxk2 was used as a loading control. Represents one of two replicates. **I.** Immunoblot was quantified relative to Hxk2 and then relative to the time of β-estradiol addition.

Using mitotically dividing cells, we further asked what would happen if both the LUTI and the PROX levels were altered. Using the same setup as above, we generated a construct with a mutated Hsf1 binding site in the *HSP60* PROX promoter (Figure 7F). Hsf1 is a transcription factor that controls the expression of genes involved in heat shock response including *HSP60* (Sakurai and Ota, 2011). We first confirmed that mutating the Hsf1-binding site reduced the basal expression of ubiGFP, but did not change LUTI transcript levels (Figure 7G-I). Upon treatment with β-estradiol, cells carrying the *hsf1* mutant construct were highly susceptible to LUTI-based repression, since Ubi-GFP abundance decreased to ∼25% of its original level after 3 hours of LUTI induction (Figure 7H and 7I). In contrast, the Ubi-GFP abundance reduced to only ∼60% of its original level in cells carrying the wild type *HSP60* PROX promoter (Figure 7H and 7I). Thus, changes in the strength of the PROX promoter also affect the extent of LUTI-based repression. Further investigation of the interplay between LUTI and PROX promoters will lead to a better understanding of when a LUTI is capable of transcriptionally repressing its PROX isoform.

## DISCUSSION

LUTIs are a recently identified class of mRNAs (Chen et al., 2017; Cheng et al., 2018; Chia et al., 2017; Van Dalfsen et al., 2018; Hollerer et al., 2019). Their 5’-extended leaders contain features, namely uORFs, that prevent the translation of the downstream ORF, and transcription of the LUTI can represses transcription from the ORF-proximal gene promoter. Consequently, LUTIs are frequently associated with a decrease in protein abundance. In this study, we show that LUTIs occur more frequently than does LUTI-based gene repression. Although we find strong evidence for translational inhibition of most LUTIs, the level of transcriptional repression is highly variable. Whether a LUTI leads to the repression of transcription from the ORF-proximal promoter is associated with the degree of H3K36me3 enrichment over the proximal gene promoter and changes to the +1 nucleosome position, and is influenced by the expression strength of both the LUTI and PROX transcripts. Our study provides the first genome-wide investigation of the key features involved in LUTI-based gene regulation.

### Developmental regulation of LUTIs

The combined use of two sequencing techniques, TL-seq and Nanopore sequencing, allowed us to determine which of the upstream TSSs give rise to full-length mRNAs, not just short intergenic transcripts (Figure 1B). Direct RNA and cDNA long-read sequencing hold immense promise for future studies of LUTIs and other transcript isoform biology (Garalde et al., 2018; Sharon et al., 2013). In addition to assisting in the identification of new splice isoforms, TSSs, and sites of transcription termination, some studies have reported accurate transcript isoform quantification by Nanopore sequencing using mixtures of spike-in RNAs (Byrne et al., 2017; Garalde et al., 2018; Oikonomopoulos et al., 2016). Unfortunately, these studies were performed with transcripts less than 2.5 kb. In our hands, a 3’ bias was observed for most transcripts, but it was strongest in transcripts greater than 2.5 kb, which includes numerous LUTIs. This 3’ bias was significant enough to prevent the accurate quantification of 5’-extended isoforms by Nanopore sequencing. The use of TL-seq for quantification allowed us to overcome this hurdle. Through the systematic identification and quantification of 5‘-extended, ORF-containing transcripts, we identified 74 genes with candidate LUTIs, which were expressed specifically in meiotic prophase.

The DNA binding protein Ume6 was enriched in the promoter of 61 LUTIs. Furthermore, the URS1 motif was conserved in 33 LUTIs across the *sensu stricto* genus (Figure 1D-H). We posit that the conservation of the regulatory binding site in such a large subset of LUTI promoters indicates that they may alter gene expression in a manner that is functionally important for meiotic progression and gamete fitness. Although Ume6 acts as a transcription repressor during vegetative growth, it interacts with the master meiotic transcription factor Ime1 in order to activate the early meiotic transcriptome (Bowdish et al., 1995). The linking of many highly conserved LUTIs to a meiosis-specific regulator inherently ties them to the progression of a developmental program: budding yeast meiosis. We predict that LUTIs will play important roles during other developmental processes because it is during these times that a cell demands dynamic and tightly regulated changes in gene expression. By employing LUTIs, a single transcription factor can both turn on and turn off gene expression in a coordinated and timely manner in order to facilitate a developmental program. In addition to the Ime1-Ume6 regulated promoters, we identified additional LUTIs that are likely regulated by other transcription factors. Since no other motifs were found, these other regulators may each play a minor role in turning on one or two LUTIs at this time in meiosis.

### uORF-mediated translational inhibition of LUTIs

uORFs are ubiquitously found in walks of life from yeast to humans. They are functionally associated with translational inhibition by sequestering the ribosomes at the 5’ end of a transcript. Various features including initiation sequence context, the distance between a uORF and the coding sequence, and uORF number in a 5’ transcript leader can all affect the degree of translational repression, but reports as to the significance of each feature vary (Calvo et al., 2009; Chew et al., 2016; Johnstone et al., 2016). What is clear is that not all uORFs are created equal. We confirm that a single uORF can lead to repression at the *NDC80* locus, but that it depends on which uORF. The uORF closest to the LUTI TSS (uORF1) does not repress translation of the *NDC80* ORF despite evidence from ribosome profiling that uORF1 is well translated. uORFs 5 and 9 both lead to robust repression even though no translation of uORF 9 is observed in meiotic cells by ribosome profiling (Figure 3C) (Brar et al., 2012). This is consistent with the genome-wide observation that greater distances between a uORF and a coding sequence correlate with greater translation efficiency of the ORF and reduced translational repression (Chew et al., 2016; Johnstone et al., 2016). In the context of the most well characterized case of uORF-mediated repression, *GCN4*, the distances between the 4 uORFs and the ORF start sequence matters greatly. That is because upon amino acid starvation, the concentration of the ternary complex, a factor required for ribosome re-initiation is decreased. This results in an extended scanning time after uORF1 before re-initiation can occur. Ultimately, the repressive uORF4 is skipped and *GCN4* is translated (Hinnebusch, 1997). Interestingly, the position of the *NDC80^LUTI^* uORF9 starts 151 bps upstream of the *NDC80* coding domain, almost exactly the same position as uORF4 with respect to the *GCN4* coding domain. Harnessing the power of yeast genetics, it could be worthwhile to further study the role of ternary complex or other trans-factors in ensuring which uORFs are integral to the translational repression of LUTIs.

The presence of translation over uORFs and a corresponding decrease in translation efficiency (TE) is a good indication that the uORFs are playing a functional role, but it does not indicate the extent of the translational regulation. Further, TE measurements at loci with LUTIs are complicated by the presence of PROX transcripts. As an alternative to measuring TE, we measured the frequency of uORF translation. When 4 or more uORFs are present in a LUTI, translation occurs only 9.4% and 20.3% of the time for the penultimate and the last uORF, respectively (Figure 3D). The high correlation between PROX transcript abundance and footprints over the ORF (Figure 2E) is also a strong indication that LUTIs provide minor, if any contribution, to ORF translation. Future use of TL-seq in combination with ribosome profiling will allow for identification of other instances in which the apparent translational regulation is due to transcript isoform toggling rather than genuine translational regulation of a single mRNA isoform. Combined with a lack of uORF selection in the 5’-extensions of LUTIs, our study has provided conclusive evidence that uORFs in LUTIs do not just dampen ORF translation, they almost entirely repress it in the vast majority of cases.

### A path towards predicting LUTI-based transcriptional repression

Previous work demonstrated that only ∼50% of the meiotically expressed 5’-extended transcript isoforms have poorly translated ORFs (Brar et al., 2012). If, as discussed above, almost all 5’-extended transcripts do not translate their ORFs, why doesn’t LUTI expression lead to translational repression more frequently? As it turns out, robust transcriptional repression of a PROX transcript by its LUTI occurs far less frequently than does translational repression of the LUTI itself. In fact, only 21 of the 72 LUTIs identified in our study have a corresponding decrease in PROX transcript to 25% of the starting abundance or less. We determined that high LUTI expression, increased H3K36me3, increased nucleosome repositioning, and longer ORFs are all significantly associated with LUTI-based transcriptional repression.

Of the features found to be important, most are associated with changes in the chromatin landscape at the PROX transcript promoter. Unexpectedly, increased H3K4me2 was not associated with LUTI-based transcriptional repression even though it is necessary for repression by *NDC80^LUTI^*(Chia et al., 2017). It is possible that only a subset of loci are dependent on H3K4me2 for repression or H3K4me2 may help to delay the kinetics of PROX re-expression later in meiosis, as it does at sites of overlapping transcription upon shifts in carbon source (Kim et al., 2017; Kim et al., 2012). Both H3K4me2 and H3K36me3 are implicated in gene repression at sites of overlapping transcription. Repression due to overlapping transcription was reported to occur on average when the TSSs were 0.9 kb apart if mediated by the Set1-H3K4me2-Set3C pathway and 2.0 kb apart when mediated by the Set2-H3K36me3 pathway (Kim et al., 2017; Woo et al., 2017). We had expected to find that LUTIs starting further from the PROX TSS would rely on H3K36me3-mediated repression and those starting closer would rely on H3K4me2. Given that H3K4me2 had no significant relationship with repression, this was not testable. However, the average distances reported above are both much longer than the 536 bp mean distance between LUTI and PROX TSSs identified in this study. This led us to hypothesize that longer distances would be associated with greater PROX downregulation, but we observed no evidence of such a relationship (ρ= −0.194, Figure 6A). It is quite possible for a LUTI to repress PROX expression differently as the distance between the two TSSs change, but in our dataset, the level of LUTI expression may mask any of those effects since it plays a far more deterministic role.

Even with features correlating significantly with repression, there were still instances in which high LUTI abundance, enrichment of H3K36me3, and changes to the +1 nucleosome were observed with no apparent repression of the PROX transcript. We hypothesized that something about the PROX promoter is different for these genes. Inspired by the hypothesis that strongly expressed PROX isoforms may be more resistant to LUTIs, we performed an in-depth analysis using a reporter construct with the *HSP60* leader to demonstrate that indeed, robustly increasing LUTI levels or decreasing PROX expression can alter the sensitivity of a gene to LUTI-based repression. This lends support to further incorporation of PROX expression level into future analyses of LUTI-based repression.

## Conclusion

In summary, we identified a group of LUTIs expressed in a coordinated manner during meiotic prophase in budding yeast. Almost all of them are severely translationally inhibited due to uORFs in their 5’-leaders. However, they do not ubiquitously lead to repression of the PROX promoters. Rather, an interplay between the strength of the promoters, the chromatin landscape, and possibly other yet to be discovered features interact to decide the transcriptional output. Future studies that integrate multiple datasets will help identify additional LUTIs and build predictive models to determine those that cause gene repression.

It still remains to be uncovered, in many cases, what the biological role of a given LUTI is and how its disruption affects cellular function. The LUTIs expressed during the endoplasmic reticulum unfolded protein response result in the downregulation of genes involved in cellular respiration, thus improving the fitness of cells under protein folding stress (Van Dalfsen et al., 2018). *NDC80^LUTI^* is essential for proper meiotic chromosome segregation (Chen et al., 2017). However, the vast majority of the LUTIs identified in budding yeast are rife for further investigation. Lastly, in human cells, the presence of a LUTI at the *MDM2* locus (Hollerer et al., 2019) opens the door for more studies into how significant LUTIs are in human development and disease.

## MATERIALS AND METHODS

### Strain Construction and Cloning

All strains for the meiotic experiments were derived from the SK1 background and contained the previously published *pCUP1-IME1/pCUP1-IME4* meiotic synchronization system (Berchowitz et al., 2013). The genotypes of all the strains used in this study are listed in Supplemental table 4. For LUTI-regulated genes of interest (*SWI4*, *ALP4*, *CDC60*, *HSP60*, and *MSC6*), the wild type gene with between 800 and 1200 bp upstream of the ORF, depending on the length of the 5’-extension, and a C-terminal 3V5 epitope tag with the *NDC80* terminator were cloned into a *LEU2* single integration vector by Gibson Assembly (Gibson et al., 2009). LUTI promoter deletion strains were similarly constructed for each gene with the cloned upstream region only containing the sequence downstream from the LUTI TSS as determined by TL-seq. In all strains, the WT copies of these 5 genes remained untouched. To construct the LexO-HSP60-ubiGFP reporter construct, a three-piece Gibson Assembly was performed in which a *HIS3* single integration plasmid carrying ubiGFP was engineered to accept fragments carrying the 8xLexO-pCYC1 and the HSP60 leader sequences. The hsf1 mutation was generated in the above plasmid by Q5 site-directed mutagenesis (E0552s, *New England Biolabs*). All single integration plasmids were digested with PmeI before transformation and the correct integrations were confirmed by PCR. All strains and plasmids used in this study are available upon request.

### pCUP1-IME1/pCUP1-IME4 Sporulation

For genome-wide cell collections, cells were prepared to progress synchronously through meiosis as described in (Chia and van Werven, 2016). Briefly, liquid YPD (1% yeast extract, 2% peptone, 2% dextrose, tryptophan (96 mg/L), uracil (24 mg/L), and adenine (12 mg/L) cultures were started and grown for ∼6 hours at 30°C until they reached an OD_600_ between 0.5 and 2.0. They were then diluted to an OD_600_ of 0.05 in reduced YPD (1% yeast extract, 2% peptone, 1% dextrose, uracil (24 mg/L, and adenine (12 mg/L) and allowed to grow for 16-18 hours at 30 °C until they reached an OD_600_ > 6. Cells were transferred to supplemented sporulation media or SPO (1% potassium acetate at pH 7.0 supplemented with adenine and uracil to 40 mg/L and histidine, leucine, and tryptophan to 20 mg/L, and 0.02% raffinose) with a final OD_600_ of 2.5 for 2 hours at 30 °C before inducing *pCUP1-IME1* and *pCUP1-IME4* with 50 µM CuSO_4_. In all other meiotic experiments, cells were prepared as in (Carlile and Amon, 2008) but with 2% potassium acetate and supplements as above. Briefly, after 24 hours of growth in YPD at RT, saturated cells (OD_600_ > 10) were diluted to an OD_600_ of 0.2-0.3 and inoculated in BYTA (1% yeast extract, 2% bacto tryptone, 1% potassium acetate, and 1.02% potassium phthalate) for 16-18 hours of growth at 30 °C (ideally to an OD_600_ of > 5). Enough cells to give a final OD_600_ of 1.85 were transferred into SPO with 2% acetate at 30 °C. After 2 hours in SPO, *IME1* and *IME4* were induced with 50 µM CuSO_4_. In meiotic experiments with the LexA-ER-AD inducible system, cells were induced with either 2 or 5 nM of β-estradiol.

### Mitotic Cell Collection

Exponentially growing cells from W303 background were back diluted to an OD_600_ of 0.2 and then treated with either 2 or 10 nM of β-estradiol. Cells were collected before induction as well as 1.5 and 3 hours after induction.

### RNA Extraction for TL-seq, Nanopore Sequencing and RNA-seq

At the indicated time points, 50 OD_600_ units of cells were collected by vacuum filtration and snap frozen in liquid nitrogen. Cells were thawed on ice and resuspended in TES buffer (10 mM Tris pH 7.5, 10 mM EDTA, 0.5% SDS) to ∼20 OD_600_. An equal volume of Acid Phenol:Chloroform:Isoamyl alcohol (125:24:1; pH 4.7) was added to cells, and they were incubated at 65°C for 45 minutes in a Thermomixer C (*Eppendorf*) shaking at 1400 RPM. The aqueous phase was transferred to a second tube of acid phenol. Samples were incubated at RT for 5 minutes while shaking at 1400 RPM in a Thermomixer. A final extraction with chloroform was performed. The aqueous phase was vortexed with chloroform for 30 seconds, separated by centrifugation, and then precipitated in isopropanol and sodium acetate for overnight at –20 °C. Pellets were washed with 80% ethanol and resuspended in DEPC water for 10 min at 37 °C. Total RNA was quantified using the Qubit RNA BR Assay Kit (Q10211, *ThermoFisher Scientific*).

### Transcript Leader Sequencing (TL-seq)

The 5’ end sequencing approach was performed as in (Wu et al., 2018). At least 5 µg of mRNA was purified from total RNA using the Poly(A)Purist MAG kit (AM1922, *Ambion*). mRNAs were fragmented for 3 minutes at 70 °C using a Zinc-based alkaline fragmentation reagent (AM8740, *Ambion*). RNAs were cleaned up using RNeasy MinElute Cleanup Kits (74204, *Qiagen*) to enrich for 200-300 nucleotide fragments. These fragments were dephosphorylated with 30 units of recombinant shrimp alkaline phosphatase (M0371, *NEW ENGLAND BIOLABS*) for 1 hour at 37 °C with RNasin Plus (N2611, *Promega*). The RNA was extracted with Acid Phenol:Chloroform:Isoamyl alcohol (125:24:1, pH 4.7) and precipitated in ethanol with 0.3 M sodium acetate and 1 µl linear acrylamide (AM9520, *Ambion*). RNA was then subjected to a decapping reaction with 2 units of Cap-Clip acid pyrophosphatase (C-CC15011H, *Tebu-Bio*) and with RNasin Plus. RNA was then again extracted using acid Phenol:chloroform:isoamyl alcohol (125:24:1) and precipitated in ethanol. Some RNA from a premeiotic time point was set apart without the decapping reaction as a non-decapping control. Subsequently, the RNA was mixed with 10 µM of custom 5’ adapter and the ligation reaction was done using T4 RNA ligase 1 (M0437M, *NEW ENGLAND BIOLABS*) and with RNasin Plus. The ligation reaction was cleaned up with the RNeasy MinElute Cleanup Kit (74204, *Qiagen*) and RNAs were mixed with 2.5 µM random hexamers (N8080127, *ThermoFisher Scientific*) and RNasin Plus, denatured at 65 °C for 5 minutes and cooled on ice. Reverse transcription reactions were carried out using SuperScript IV reverse transcriptase (18090010, *Invitrogen*). The RNA templates were degraded by incubating reactions with 5 units of RNase H (M0297, *NEW ENGLAND BIOLABS*) and 1.0 µl of RNase cocktail enzyme mix (AM2286, *Ambion*). DNA products were purified using 1.8x volume of HighPrep PCR beads (AC-60050, *MagBio*). Purified products were subjected to second strand synthesis using 0.3 µM of second strand biotinylated primer and the KAPA Hi-Fi hot start ready mix (KK2601, *Roche*). The second strand reaction was carried out at 95 °C for 3 minutes, 98 °C for 15 seconds, 50 °C for 2 minutes, 65 °C for 15 minutes and held at 4 °C. Double stranded product (dsDNA) was purified with 1.8x volume HighPrep PCR beads and concentration was quantified using the Qubit dsDNA HS assay kit (Q32851, *Invitrogen*). 25 ng of dsDNA was then used as input for the KAPA Hyper Prep Kit (KK8504, *Roche*) and ligated to KAPA single indexed adapters Set B (KK8702, *Roche*). Samples were processed according to manufacturer’s instructions with one exception: just prior to the library amplification step, samples were bound to MyOne Streptavidin C1 Dynabeads (65001, *ThermoFisher Scientific*) to capture biotinylated dsDNA. Library amplification over 14 PCR cycles was done on the biotinylated dsDNA fraction bound to the beads. Amplified libraries were quantified by Qubit, and adapter-dimers were removed by electrophoresis of the libraries on Novex 6% TBE gels (EC62655BOX, *Invitrogen*) at 120 V for 1 hour, and excising the smear above ∼150 bp. Gel slices containing libraries were shredded by centrifugation at 13000 *g* for 3 minutes. Gel shreds were re-suspended in 500 µl crush and soak buffer (500 mM NaCl, 1.0 mM EDTA and 0.05% v/v SDS) and incubated at 65 °C for 2 hours on a thermomixer (1400 rpm for 15 seconds, rest for 45 seconds). Subsequently, the buffer was transferred into a Costar SpinX column (8161, *Corning Incorporated*) with two 1 cm glass pre-filters (1823010, *Whatman*). Columns were centrifuged at 13000 *g* for 1 minute. DNA libraries in the flowthrough were precipitated at −20 °C overnight in ethanol with 0.3 M sodium acetate and 1 µl linear acrylamide (AM9520, *Ambion*). Purified libraries were further quantified and inspected on a Tapestation (*Agilent Technologies, Inc*). The libraries were sent for 100 bp SE sequencing on an Illumina HiSeq 4000 at the Vincent J. Coates Genomics Sequencing Laboratory.

From the sequencing reads, the 3’ Illumina adapter (AGATCGGAAGAGC) was trimmed using cutadapt with the --*minimum-length* option set to 20 bp (v2.3, (Martin, 2011)). From the 3’ trimmed output, the 5’ Illumina adapter (CACTCTGAGCAATACC) was trimmed from reads by cutadapt. To select for reads with the most 5’ end of a transcript, only reads in which the 5’ adapter was recognized and then trimmed were carried forward. Reads were aligned by STAR (v2.5.3a, (Dobin et al., 2013)) using indices generated from an SK1 genome assembled by combined PacBio and Illumina sequencing (Yue et al., 2017). A custom SK1 genome was forged with BSgenome (v1.50.0, Pagè, 2018) using the above assembly (Yue et al., 2017). Bam files were imported into CAGEr and the CAGEr pipeline was applied to define TSSs and quantify transcript abundances as follows (v1.24.0, Haberle et al., 2015). Reads at TSSs were counted (*getCTSS*) and normalized by *“simpleTpm”* (*normalizeTagCount*). An initial clustering was performed (*clusterCTSS* with *threshold = 2*, *thresholdIsTpm =TRUE*, *method = “distclu”*, *maxDist = 5*, *removeSingletons = TRUE*, and *keepSingletonsAbove = 3*), and the output was aggregated into larger clusters representative of all the activity expected from a single promoter (*aggregateTagClusters* with *tpmThreshold = 1* and *maxDist = 50*). Clustered TSSs were exported as bedGraph files for visualization in IGV (*exportCTSStoBedGraph* with *values = “normalized”*). Cluster counts were exported to DESeq2 by time point (*concensusClustersDESeq2*), and fold-changes were calculated by DESeq2 with default settings. Output from this clustering was used to define TSSs coordinates of 5’-extended transcripts in the pipeline below. A secondary and more permissive clustering (*threshold = 1, thresholdIsTpm =TRUE, method = “distclu”, maxDist = 5, removeSingletons = FALSE*) was performed after LUTIs were defined. Output from the secondary clustering was used for quantification and in all presented TL-seq scatterplots. The TL-seq used to define LUTI promoters was performed in triplicate. All TL-seq comparisons to RNA-seq were performed in duplicate.

### polyA Selection for Nanopore Sequencing and RNA-seq

PolyA selection was performed on 100 µg of RNA using 150 µL of oligo-dT DynaBeads (61002, *ThermoFisher Scientific*). RNA was denatured at 80 °C for 2 minutes in binding buffer (10 mM Tris-HCl pH 7.5, 0.5 M LiCl, 3.35 mM EDTA) before being placed on ice. At RT the oligo-dT beads were added to the sample and together they were incubated at room temperature of 5 minutes. Beads were washed 2x in Buffer B (10 mM Tris-HCl pH 7.5, 0.15 M LiCl, 1 mM EDTA). PolyA-selected RNA was eluted from the beads by heating at 80 °C for 2 minutes in 10 mM Tris 7.0. It was quantified with a Qubit using the RNA HS assay kit (Q32852, *ThermoFisher Scientific*).

### Nanopore Sequencing

500 ng of polyA-selected RNA was used as directed in the Direct RNA Sequencing Kit (SQK-RNA001, *Oxford Nanopore Technologies*). The library was loaded onto a minION (MIN-101B, *Oxford Nanopore Technologies*) with an R9.4.1 flow cell (FLO-MIN106, *Oxford Nanopore Technologies*). MinKNOW (v1.10.23, *Oxford Nanopore Technologies*) was run without live base calling for 48 hours.

Bases were called from fast5 files with the Albacore script read_fast5_basecaller.py (v2.1.10, *Oxford Nanopore Technologies*). 491,142 reads were sequenced. Reads were aligned to the genome with minimap2 (v2.9-r720, (Li, 2018)) using options *–ax splice – k14 –uf*. Bam files were visualized directly in IGV.

### Pipeline for 5’-extended Transcript Discovery

Using the output from DESeq2 after CAGEr, TSS clusters were filtered for coordinates in which the mean over both time points was > 2 transcripts per million and the log2 fold-change as cells entered meiotic prophase compared to premeiotic was > 2. After applying these filters, the coordinates for each peak were manually inputted into IGV. The TL-seq peak was compared to Nanopore sequencing reads from a sample taken during meiotic prophase (4 hours). If at least one Nanopore read extended from a region near the TSS coordinates and continued uninterrupted across the entirety of a neighboring ORF, the coordinates were marked for continued investigation. Purely intergenic and either 5’ or 3’ truncated transcripts were removed in this way. From the remaining subset of peaks, a 5’-extension was only called if a second promoter, downstream, but on the same strand, was closer to the ORF. Through this criterion, canonical meiosis-specific genes were eliminated from the analysis. It resulted in 74 candidate LUTIs with 5’-extensions. For downstream analyses, the single most dominant bp in each TSS cluster was determined by a custom python script.

### Motif Discovery

Meme was applied to the sequences 300 bp up and 300 bp downstream from each LUTI TSS with options –*w 10 –dna –revcomp.*(Bailey and Elkan, 1994). A motif was considered significant in an individual sequence if it had a combined match p-value < 0.05.

### Chromatin Immunoprecipitation (ChIP)

For Ume6-3V5 ChIP, 300 OD_600_ units of stationary phase cells in BYTA. In histone modification experiments, 112.5 OD_600_ units of cells were collected during both premeiotic stage and meiotic prophase. In all instances, cells were fixed with 1% formaldehyde. The formaldehyde was quenched with 125 mM glycine, cells were pelleted, washed with PBS, and then lysed by Beadbeater (Mini-Beadbeater-96, *Biospec Products*) with zirconia beads 4 x 5 minutes in FA Buffer (50 mM Hepes pH 7.5, 150 mM NaCl, 1 mM EDTA, 1% Triton, 0.1% sodium deoxycholate, 0.1% SDS supplemented with cOmplete protease inhibitor tablets (11873580001, *Roche*). Note that for the Ume6-3V5 ChIPs, due to the number of cells collected, lysates were prepared in 3 separate tubes. They were processed separately until after the IP. Lysates were collected and centrifuged for 3 minutes at 2000 g. The supernatants were transferred to fresh tubes and centrifuged for 15 minutes at 20,000 g. The supernatant was discarded, and the pellet of chromatin was resuspended in 1 mL FA Buffer. Samples were sonicated: 30 seconds on, 30 second off, for 5 minutes with a Bioruptor Pico (*Diagenode*) to an average fragment size of ∼ 200 bp. The supernatant from a 1 minute 20,000 g centrifugation was carried forward to the IP.

From the isolated chromatin, 30 µL were set aside as input. For Ume6-3V5 ChIPs, to each of 3, 600 µL chromatin aliquots, 1 uL of a mouse α-V5 antibody (R960-25, *Thermo Fisher*) was added. For histone modification ChIPs, 3 µL of an antibody specific to either H3K36me3 (ab9050, *abcam*) or H3K4me2 (ab32356, abcam) was added to the chromatin. Samples were incubated for 2 hours with nutation. Protein A Dynabeads (10001D, *Invitrogen*) were blocked with 0.1% BSA in FA Buffer for at least 2 hours at 4°C. They were washed 2 x with FA Buffer, resuspended with FA Buffer to their original volume, and 10-20 µL (10 for V5 and 20 for histone modification IP) of the resuspended beads were added to each tube of chromatin. The chromatin-bead mixture was nutated at 4 °C overnight. The IP was washed 6 times: 2x FA Buffer, 2x Buffer 1 (FA Buffer, 260 mM NaCl), and 2x Buffer 2 (10 mM Tris pH 8.0, 250 mM LiCl, 0.5% NP-40, 0.5% sodium deoxycholate, 1 mM EDTA). Between each wash, samples were nutated for 5 minutes at 4 °C. To IP and input samples, 150 µL or 120 uL of TE (10 mM Tris pH 8.0, 1 mM EDTA) with 1% SDS was added, respectively. The precipitate was eluted from the beads by shaking at 1200 RPM in a Thermomixer (*ThermoFisher Scientific*) at 65 °C. Eluates were treated with ∼0.33 mg/mL RNase A (12091021, *Invitrogen*) for 30 minutes at 37 °C and then with ∼1.2 mg/mL Proteinase K (3115879001, *Roche*) overnight at 65°C. Samples were cleaned up with Qiagen QiaQuick PCR Purification Kit (28106, *Qiagen*) and eluted in EB. DNA was quantified by Qubit with the dsDNA HS Assay Kit (Q32854, *Invitrogen*). Libraries were prepared as instructed by the ThruPLEX DNA-seq Kit (R400427, *Takara*). For input and H3K36me3 IP samples, 15 ng of starting material was used with 7 rounds of PCR. For all other IP samples, 0.5-1 ng of starting material was used with 11 rounds of PCR. AMPure XP beads (A63881, *Beckman Coulter*) were used to select fragments between 200-500 bp. Samples were submitted for 50 bp SE sequencing by the Vincent J. Coates Genomics Sequencing Laboratory with a HiSeq4000.

### Ume6 ChIP-seq Analysis

Reads were aligned to the SK1 genome with bowtie2 (v2.3.4.3, Langmead and Salzberg, 2012; Langmead et al., 2019). Using randsample from macs2, all libraries were down-sampled to 2 million reads (v2.1.1.20160309, Zhang et al., 2008). Macs2 callpeak was used to call peaks in IP samples over input samples with options *–B –q 0.001 --keep-dup “all” --call-summits –nomodel –extsize 147*. Bigwig files for viewing in IGV were generated by macs2 bdgcmp with option *–m FE* followed by bedGraphToBigWig (v4, (Kent et al., 2010)). Heatmaps and metagene plots centered around TSSs as defined by TL-seq were constructed with deeptools2 (v3.0.1, (Ramírez et al., 2016)). A 5’-extended or canonical target promoter was considered to be enriched by Ume6 if, in at least 2 of 3 ChIP replicates, a peak was called (log2 fold-change > 2 over input) within 300 bp of the transcript’s TSS.

### Conservation of URS1 Binding Sites

The 5’-extended and canonical Ume6 targets enriched with both Ume6 and a URS1 binding site in their promoters were analyzed for degree of conservation. Meme was run on the sequences +/- 300 bp from their TSSs with options *–w 10 –dna –revcomp* was run to identify the location of the URS1 binding sites with regard to the TSS. Using a custom python script, the midpoint of each URS1 binding motif was determined relative to the ORF associated with either the LUTI or canonical target. Subsequently, the chromosome locations of the URS1 midpoint in the sacCer3 S288C reference genome were found. To assess conservation of the regions around URS1 motifs at 5’-extended and canonical targets, phastCons (Siepel et al., 2005) was performed with options *– target-coverage 0.025-expected-length 12 –rho 0.4*. The tree phylogeny model and the genome alignments of *S. cerevisiae, S. paradoxus, S. mikatae, S. kudriavzevii, S. bayanus, S. castelli, and S. kluyveri* were from (Siepel et al., 2005). However, the alignments of *S. castelli* and *S. kluyveri* were excluded in all analyses here because these yeast species have lost the Ime1 and Ume6 genes. Metagene plots and heatmaps of conservation were generated with deeptools2 (v3.0.1, Ramírez et al., 2016)

### RNA-seq

The RNA-seq libraries were prepared using the NEXTflex^TM^ Rapid Directional mRNA-Seq Kit (NOVA-5138, *Bioo Scientific*) according to manufacturer’s instructions. 100 ng of poly-A selected mRNA was used for all libraries. Libraries were quantified using the Agilent 4200 TapeStation (*Agilent Technologies, Inc.).* AMPure XP beads (A63881, *Beckman Coulter*) were used to select fragments between 200-500 bp. Samples were submitted for 150 bp SE sequencing by the Vincent J. Coates Genomics Sequencing Laboratory with a HiSeq4000.

Quantification of RNA as transcripts per million was done using salmon in the mapping-based mode with mapping validation (v0.13.1, Patro et al., 2017). Fold-change quantification was performed by DESeq2 with counts generated by summarizeOverlaps using default options (v1.22.2, Love et al., 2014). Scatterplots were generated with matplotlib (Hunter, 2007).

### uORF Analysis

ATGs were counted and the codon frequency was determined with a custom python script. For genes with LUTIs, the counts and codon frequencies were determined for the region between the PROX TSS and the LUTI TSS. For all other genes, sequences from the 500 bp upstream of the TSS were used.

LUTIs with > 4 uORFs were analyzed to determine which of the uORFs were translated. Footprints were quantified for the first 6 codons of each uORF using the tools in (Brar et al., 2012) and (Ingolia et al., 2009). The ribosome footprinting data was taken from the 3 hr timepoint in (Cheng et al., 2018). Any uORFs with at least 4 footprint reads found across the first 6 codons of the gene were considered to be translated.

### Immunoblotting

Immunoblotting was performed as previously described in (Chen et al., 2017). To track tagged proteins of interest, membranes were incubated with either a mouse α-V5 antibody (R960-25, *Thermo Fisher*) or a mouse α-GFP antibody (632381, *Takara*) to a dilution of 1:2000 in Odyssey Blocking Buffer (PBS) (*LI-COR Biosciences*) with 0.01% Tween. Both V5 and GFP immunoblots were also incubated with a rabbit α-hexokinase (α-Hxk2) antibody (H2035, *US Biological*) diluted between 1:15,00-1:20,000. Secondary antibodies included a α-mouse antibody conjugated to IRDye 800CW (926-32212, *LI-COR Biosciences*) and a α-rabbit antibody conjugated to IRDye 680RD (926-68071, *LI-COR Biosciences*). They were each diluted to 1:15,000 in Odyssey Blocking Buffer (PBS) with 0.01% Tween. An Odyssey system (*LI-COR Biosciences*) was used to image all blots, and Image Studio Lite (*LI-COR Biosciences*) was used for all quantification.

### RNA Extraction for Blotting and RT-qPCR

At the indicated time points, between 0.4 and 5 OD_600_ units of cells were collected by centrifugation and snap frozen in liquid nitrogen. Cells were thawed on ice and resuspended in TES (10 mM Tris pH 7.5, 10 mM EDTA, 0.5% SDS). An equal volume of Acid Phenol:Chloroform:Isoamyl alcohol (125:24:1; pH 4.7) was added to cells, and they were incubated at 65°C for 45 minutes in a Thermomixer C (*Eppendorf*) shaking at 1400 RPM. The aqueous phase was transferred to a second tube with chloroform. The aqueous phase was vortexed with the chloroform for 30 seconds, separated by centrifugation, and then precipitated in isopropanol and sodium acetate overnight at –20°C. Pellets were washed with 80% ethanol and resuspended in DEPC water for 10 min at 37 °C. Total RNA was quantified by Nanodrop.

### RNA Blotting

RNA blot analysis protocol was performed as described previously (Koster et al., 2014) with minor modifications. 8 µg of total RNA was denatured in a glyoxal/DMSO mix (1M deionized glyoxal, 50% v/v DMSO, 10 mM sodium phosphate (NaPi) buffer pH 6.5–6.8) at 70 °C for 10 minutes. Denatured samples were mixed with loading buffer (10% v/v glycerol, 2 mM NaPi buffer pH 6.5–6.8, 0.4% w/v bromophenol blue) and separated on an agarose gel (1.1% w/v agarose, 0.01 M NaPi buffer) for 3 hr at 100 V. The gels were then soaked for 25 minutes in denaturation buffer (0.05 N NaOH, 0.15 M NaCl) followed by 20 minutes in neutralization buffer (0.1 M Tris-HCl pH 7.5, 0.15 M NaCl). RNA was transferred to nitrocellulose membrane for 1 hour via vacuum transfer as described in Stratagene’s Membranes Instruction Manual (*Stratagene*) and crosslinked using a Stratalinker UC Crosslinker (*Stratagene*). rRNA bands were visualized using methylene blue staining. The membranes were blocked in ULTRAhyb Ultrasensitive Hybridization Buffer (AM8669, *ThermoFisher Scientific)* for 3 hours before overnight hybridization. Membranes were washed twice in Low Stringency Buffer (2x SSC, 0.1% SDS) and three times in High Stringency Buffer (0.1X SSC, 0.1% SDS). All hybridization and wash steps were done at 42 °C. Radioactive probes were synthesized using a Prime-It II Random Primer Labeling Kit (300385, *Agilent Technologies, Inc*).

### Chromatin modification analysis

Reads were aligned to the SK1 PacBio genome assembly with bowtie2 (v2.3.4.3, Langmead and Salzberg, 2012). Macs2 callpeak was used to call peaks in IP samples over input samples with options *–B –q 0.01 –nomodel –extsize 147*. Bigwig files for viewing in IGV and for further quantification were generated by macs2 bdgcmp with option *–m FE* followed by bedGraphToBigWig (v4, Kent et al., 2010; Zhang et al., 2008). To quantify the change in H3K36me3 and H3K4me2 enrichment over the promoters of PROX transcripts, fold enrichment scores were extracted from regions 50 bp upstream and 500 bp downstream from the PROX TSS with bedtools (Quinlan and Hall, 2010). With custom python scripts, the scores from each bp of the upstream regions and each bp of the downstream regions were summed for each gene. The ratio of the upstream and the downstream region enrichments were quantified and the change in the score from premeiotic stage to meiotic prophase was determined. Ultimately, the mean of the fold-change was calculated from samples in triplicate. Heatmaps and metagene plots were prepared with deeptools2 (Ramírez et al., 2016).

### Micrococcal Nuclease Sequencing (MNase-seq)

The protocol was adapted from Basic Protocol 1 in (Rodriguez et al., 2014) with the following changes. In premeiotic stage and meiotic prophase, 112.5 OD_600_ units of cells were fixed in 1% formaldehyde with light shaking at RT for 15 minutes. Crosslinking was quenched by 125 mM of glycine for 5 minutes at RT. Cells were pelleted and washed twice with ice cold milliQ water. Cells were spheroplasted in 20 mL of Spheroplast Solution (1 M Sorbitol, 50 mM Tris pH 7.5, 10 mM β-ME) with 100 µL of 10 mg/mL zymolase until they appeared non-refractive and shadow-like after ∼20-30 minutes. Spheroplasted cells were resuspended in 2 mL MNase Digestion Buffer (1 M Sorbitol, 50 mM NaCl, 10 mM Tris pH 7.5, 5 mM MgCl_2_, 1 mM CaCl_2_, 0.075% NP-40, 0.5 mM spermidine, 1 mM β-ME) if collected during premeiotic stage and 4 mL if collected during meiotic prophase, after completion of S-phase. Digestions were performed with 600 uL of spheroplasts, 30 units of Exonuclease III (M0206S, *New England Biolabs*), and either 10, 20, or 40 units of MNase (LS004797, *Worthington*). Crosslinks were reversed, protein was degraded by Proteinase K (3115879001, *Roche*), and a phenol/chloroform/isoamyl alcohol DNA extraction, ethanol precipitation, RNase A (12091021, *Invitrogen*) treatment, and phosphatase treatment were performed as described previously (Rodriguez et al., 2014). Size selection was performed by running samples on a 1.8% LMT agarose gel at 80 V for 40 minutes at room temperature and gel extracting the mononucleosome band with a Monarch Gel Extraction Kit (T1020S, *New England Biolabs*). Note that of the samples digested with 10, 20, and 40 units of MNase, only the samples with a ratio of mononucleosomes to dinucleosomes closest to 80/20 were size selected and carried forward for library preparation. Gel extracted samples were quantified by Qubit with the dsDNA HS Assay Kit (Q32854, *Invitrogen*). Libraries were prepared with 50 ng starting material as instructed by the ThruPLEX DNA-seq kit (R400427, *Takara*). Amplification was performed with 5 rounds of PCR. AMPure XP beads (A63881, *Beckman Coulter*) were used to select fragments between 150-500 bp. Samples were submitted for 100 bp PE sequencing to the Vincent J. Coates Genomics Sequencing Laboratory with a HiSeq4000.

Reads were aligned to the SK1 genome with bowtie2 (v2.3.4.3, Langmead and Salzberg, 2012). To select for only fragments between 130 and 170 bps, alignmentSieve from deeptools2 was performed (Ramírez et al., 2016). BigWig files were generated by bamCoverage with options -*-MNase –bs 1 --normalizeUsing CPM* (Ramírez et al., 2016). DANPOS (v2.2.2) was run to determine various aspects of nucleosome location and occupancy and fuzziness (K. Chen et al., 2015, 2013). A custom python script was used to assign the locations of +1 and –1 nucleosome with respect to PROX TSSs.

### Reverse Transcription-Quantitative Polymerase Chain Reaction (RT-qPCR)

To 5 μg of isolated total RNA in 1x DNase I reaction buffer, 1 unit of DNase I was added. The reaction was incubated at 37 °C for 30 minutes. The DNase-treated RNA was extracted by Acid Phenol:Chloroform:Isoamyl alcohol (125:24:1; pH 4.7) and precipitated with isopropanol and sodium acetate overnight. The RNA was washed in 80% ethanol and resuspended in milliQ water. cDNA was reverse transcribed following the Superscript III kit (18080044, *ThermoFisher Scientific*). Quantification was performed with Absolute Blue qPCR Mix (AB4162B, *ThermoFisher Scientific*). The primers used to quantify *ubiGFP^LUTI^* included a forward primer which annealed immediately upstream of the *HSP60^PROX^*TSS and a reverse primer which annealed to the first bases of ubiquitin. Meiotic signals were normalized to *PFY1* and mitotic signals were normalized to *ACT1*. Oligonucleotides are in Supplemental Table 5.

## Data availability

Data generated in this study are available at NCBI GEO under the accession ID GSE140177 (https://www.ncbi.nlm.nih.gov/geo/query/acc.cgi?acc=GSE140177). The custom code used for the analysis is available in the following code repository: https://github.com/atresen/LUTI_key_features.

## ACKNOWLEDGEMENTS

We thank Nicholas Ingolia and Gloria Brar for their help on the analysis of our TL-seq data alongside ribosome profiling, and Cai Li for his help in determining how to quantify the TL-seq data. We thank Michael Eisen, Nicholas Ingolia, Gloria Brar, Jasper Rine, Jingxun Chen, Tina Sing, Kate Morse, Amanda Su, Eric Sawyer, Grant King and Andrea Higdon for their experimental suggestions and/or comments on this manuscript as well as all the members of the Ünal and Brar labs for their input on the project. This work is supported by funds from the Pew Charitable Trusts (00027344), Damon Runyon Cancer Research Foundation (35-15) and National Institutes of Health (DP2 AG055946-01) to EÜ.

## AUTHOR CONTRIBUTIONS

AT: conceptualization, data curation, formal analysis, investigation, methodology, validation, visualization, and drafted the manuscript. VJ: data curation, formal analysis, investigation, methodology, validation, and visualization, MC: performed the TL-seq library preparations and manuscript editing. HL: investigation for the *NDC80* uORF analysis. FJW: supervision, project administration and manuscript editing. EÜ: conceptualization, formal analysis, methodology, visualization, supervision, project administration, funding acquisition, and drafted the manuscript.

